# The historical biogeography of range expansion predicts spatial patterns in reproductive assurance

**DOI:** 10.1101/2024.11.04.621881

**Authors:** Mackenzie Urquhart-Cronish, Ken A. Thompson, Amy L. Angert

## Abstract

Plant reproductive assurance describes the ability of a plant to successfully reproduce in an environment that is potentially devoid of conspecifics and/or pollinators. Traditionally, studies have focused on the role of contemporary ecology—-such as pollinator or mate availability—-in driving spatial patterns in reproductive assurance within species, however, historical processes such as post-glacial range expansion may be an understudied alternative explanation for geographic variation in mating system. This is because during the process of geographic range expansion into novel habitat, selection should favour individuals that possess traits promoting reproductive assurance (i.e., autonomous selfing or clonal reproduction). Here, we used Northern pink monkeyflower—*Erythranthe* (*Mimulus lewisii*—a hermaphroditic, self-compatible, perennial, alpine plant, as a focal species to investigate the historical signatures of geographic range expansion by combining phylogeographic analyses with a greenhouse survey of range-wide reproductive assurance. First, we detected significant geographic variation among populations in two components of reproductive assurance: self-fertilization and clonal propagation. Next, using genome-wide single nucleotide polymorphism (SNP) data, we identified three distinct genetic clusters structured by longitude and estimated geographic coordinates of the most likely origins of range expansion within each cluster. We found that both measures of reproductive assurance significantly increased on average with distance from the inferred biogeographic origin, which is consistent with the hypothesis that reproductive assurance undergoes adaptive evolution during range expansion. This study supports hypotheses underlying Baker’s Law and contributes to our understanding of spatial variation in reproductive assurance, which is linked to variation in evolutionary potential, adaptability, and the long-term persistence of populations across large spatial scales.

**Lay summary:** What generates and maintains the amazing diversity in plant reproduction we see today? Traditionally, studies have focused on the role of contemporary ecology, such as pollinator or mate availability, in driving spatial patterns within species, however, historical drivers may be an understudied alternative explanation. Here, we test the hypothesis that expansion into new habitats after glacial retreat, where pollinators or mates were scarce, favors plants with the ability to reproduce in the absence of pollinators and mates by making self-pollinated seeds or vegetative clones of themselves. We find support for the hypothesis, where the probabilities of making both autonomously selfed seed and clonal vegetative propagules increase on average along inferred pathways of range expansion. Our study joins a handful of previous studies that tested and provided support for historical range expansion shaping range-wide spatial variation in reproductive assurance in plants, suggesting post-glacial range expansion may be a common mechanism creating diversity in plant mating systems.

## Introduction

Patterns of who mates with whom in natural populations have important implications for genetic variation, adaptability, and individual fitness (Whitehead et al., 2018). Although outcrossing can maintain higher levels of genetic diversity—and therefore greater evolutionary potential— many populations have independently evolved from predominantly outcrossing to partial or even complete self-fertilization (Schemske and Lande, 1985). This transition can be explained by the “reproductive assurance” hypothesis, wherein the ability to self-fertilize or propagate clonally provides a mechanism to successfully reproduce in novel environments potentially devoid of conspecifics and/or suitable pollinators. Studies of the reproductive assurance hypothesis often focus on how contemporary ecology, like pollinator or conspecific densities, can drive spatial patterns in plant reproductive diversity (Fausto et al., 2001; Ghazoul, 2005; Hargreaves and Eckert, 2014; Kalisz et al., 2004; Vaughton and Ramsey, 2010).

On contemporary timescales, demographic factors—such as low population density—can influence ecological interactions that drive the evolution of autonomous selfing. This is because an individual that has the ability to mate with itself will experience a fitness advantage in a low-density population, where conspecific mates are scarce and pollinators are less abundant and effective (Kalisz et al., 2004). In contrast, in a high-density population, an individual with a high level of self-fertilization could experience a fitness disadvantage, due to the higher degree of inbreeding depression expected among its offspring (Griffin and Willi, 2014). Therefore, geographic variation in self-compatibility can be driven by current variation in demography and pollinator abundances across a species’ range (Kalisz et al., 2004; Opedal et al., 2016).

Reproductive assurance can also be achieved via asexual reproduction, such as clonal growth (Baker, 1955; Pannell et al., 2015), which is the most common form of vegetative propagation in angiosperms (Barrett, 2015). Clonal growth occurs through a diverse array of plant organs (e.g., bulbs, rhizomes, and stolons) that vary in their dispersal capabilities from the parent plant. Clonal reproduction is commonly associated with perenniality, longevity, and occurrence in habitats in which sexual recruitment is often restricted, such as the colonization of new habitat (Baker, 1955; Pannell et al., 2015). Generally, clonality reduces population genetic diversity and can increase inbreeding when genets have more than one flower open simultaneously (i.e., geitonogamy) (Charpentier, 2001; Handel, 1985; Vallejo-Maŕın et al., 2010).

However, contemporary ecological explanations for spatial variation in plant reproductive diversity are not always supported (Busch and Delph, 2012; Herlihy and Eckert, 2005; Koski et al., 2017), suggesting alternative, understudied, factors might influence range-wide plant reproductive diversity. An alternative explanation for spatial variation in reproductive assurance across a species’ geographic range is the role of post-glacial range expansion (Baker, 1955; Koski et al., 2019; Pujol et al., 2009). This idea stems from a classic observation made by Herbert Baker, who observed that the successful colonization of a novel habitat is facilitated by the ability of a species’ to self-fertilize and/or reproduce clonally (Baker, 1955; Cheptou, 2012). This highlights how demography during historical colonization events (e.g., population bottlenecks, low-population density) can drive the evolution of autonomous selfing and/or clonal propagation, whereby successful reproduction can occur in the absence of pollinator visitation. Importantly, it should be noted that both contemporary and historical hypotheses of drivers of reproductive assurance could be in effect simultaneously. Compared to the role of contemporary ecology, the role of historical range expansion has been relatively understudied because high-caliber tests require data on both range-wide variation in reproductive assurance and phylogeographic reconstructions (i.e., genomic data). With increasing access to large genomic datasets, a handful of studies have tested these questions and found support for increased autonomous selfing ability with increasing distance from inferred refugia (Koski et al., 2019; Willi et al., 2018; Zeitler et al., 2023). Using the native plant *Erythranthe* (*Mimulus*) *lewisii* as our focal species, we test the hypothesis that serial founder events during historical range expansions drive the selection for reproductive assurance towards the range edge. Based on the potential for multiple historical refugia to contribute to range expansion following continental glacier retreat (Hewitt, 2000; Shafer et al., 2010), we first define putative independent range expansion events by investigating range wide spatial population genetic structure. Given the resolved spatial genetic structure, more specifically we predict that *(i)* there will be geographic variation in reproductive assurance, as measured by autonomous selfing ability and clonal reproduction and *(ii)* in the context of historical range expansion, populations further from historical refugia will have greater ability to self-fertilize and produce clonal rhizomes.

## Methods

### Study system

*Erythranthe (Mimulus) lewisii* (Phrymaceae), Northern pink-monkeyflower, is an herbaceous perennial plant native to subalpine and alpine riparian habitat throughout Northwestern North America. It is pollinated primarily by bumblebees, but has hermaphroditic flowers and is capable of self-fertilization both within (likely via delayed selfing during corolla abscission [Dole 1992]) and between flowers, and is capable of reproducing clonally via rhizomes. Seed dispersal in monkeyflowers is thought to primarily occur via water (Waser et al., 1982), but dispersal can also potentially occur via wind and mammals (Vickery et al., 1986). Here, we focus on *E. lewisii* across its entire geographic range (Fig. 1).

**Figure 1:**
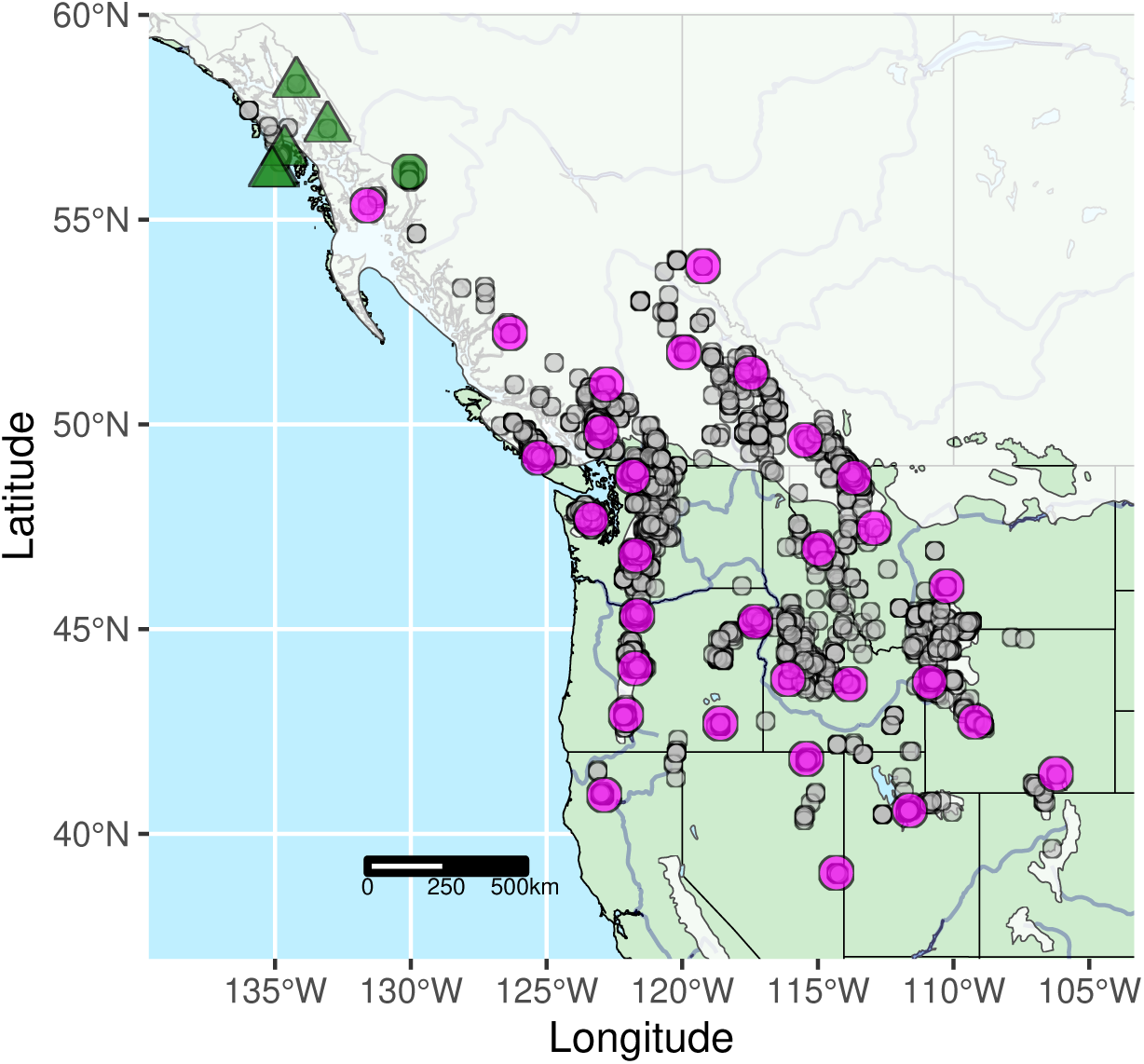
Map of range wide occurrence records for *Erythranthe lewisii* (grey points) with sampling locations (magenta and green points) and last glacial maxima of the Cordilleran ice sheet (white overlay) (Dalton et al., 2020). Grey points are filtered range wide occurrence records (collected from GBIF, Alaska Center for Conservation Science, and BEC Data Base), magenta points are sites with seed and leaf tissue collections, green points are sites with only leaf tissue collections, circles are field-collected samples (n = ∼10-20/site), and triangles are herbarium samples (n = 1 per site). See Figure S1 for decomposed maps of sampling vs. occurrence records and sampling vs. continental ice sheet.

### Range wide sampling

We conducted fieldwork between August and September 2019 (one site), 2020 (five sites), 2021 (21 sites) and 2022 (four sites) where we sampled leaf tissue or mature seed & leaf tissue from a total of 31 sites spanning the geographic range of *E. lewisii* (Table 1; Fig. 1). We sampled leaf tissue and seed from ∼ 10–20 plants per population, with a distance of at least ∼ 1 m between the samples to avoid sampling clonal ramets of the same genet. We collected leaf tissue and seed samples from a single plant separately in coin envelopes stored on non-toxic indicator desiccant (Dry & Dry Orange Indicating Silica Gel Desiccant Beads). We supplemented leaf tissue collections from the northwestern distribution limit (Alaskan Panhandle) with leaf tissue from five herbarium specimens (see Supp Mat for accession details) (Fig. 1).

**Table 1:** Summary of sampling sites, sample sizes, and population-level genetic diversity stats organized by ancestral cluster and increasing distance from putative origin of expansion (within ancestral genetic cluster, see main text for locations of refugia). Note, (*) AK is made up of three herbarium samples that passed genomic quality filtering (56.5388, -134.7578; 56.4162; -134.8367; 57.225, -133.07) and were merged to create a single experimental population. Two populations (RW15, RW18) were detected as having hybrid ancestry from ADMIXTURE analysis, as indicated in the “Cluster” column.

| Site | $N_{\text{genomic}}$ | $N_{\text{gh}}$ | Province/<br>State | Latitude | Longitude | Elevation (m) | Cluster | Distance from<br>refugia (km) | He | Ho | PA count | $\pi$ | $F_{IS}$ |
| --- | --- | --- | --- | --- | --- | --- | --- | --- | --- | --- | --- | --- | --- |
| RW01 | 11 | 7 | BC | 49.19 | -125.29 | 1169 | Western | 30.69 | 0.04 | 0.02 | 482.32 | 0.04 | 0.92 |
| RW03 | 11 | 1 | WA | 47.68 | -123.35 | 1170 | Western | 235.11 | 0.05 | 0.03 | 701.80 | 0.05 | 0.90 |
| RW35 | 12 | 9 | BC | 52.23 | -126.35 | 963 | Western | 383.04 | 0.04 | 0.02 | 259.97 | 0.04 | 0.92 |
| RW33 | 9. | 3 | AK | 55.35 | -131.59 | 812 | Western | 828.93 | 0.07 | 0.01 | 186.98 | 0.07 | 0.94 |
| RW10 | 12 | – | BC | 56.17 | -130.05 | 1142 | Western | 887.77 | 0.04 | 0.01 | 249.19 | 0.04 | 0.94 |
| RW32 | 11 | 10 | CA | 40.95 | -122.87 | 2019 | Western | 950.52 | 0.03 | 0.02 | 1761.86 | 0.04 | 0.94 |
| AK* | 3 | – | AK | 57.23 | -133.07 | 1353 | Western | 1089.37 | 0.06 | 0.02 | 0.95 | 0.07 | 0.93 |
| RW31 | 12 | 4 | WA | 48.73 | -121.83 | 1571 | Central | 139.55 | 0.05 | 0.04 | 308.13 | 0.06 | 0.60 |
| RW02 | 11 | 8 | WA | 46.80 | -121.71 | 1560 | Central | 170.67 | 0.06 | 0.04 | 459.70 | 0.06 | 0.60 |
| RW30 | 12 | 6 | BC | 49.81 | -122.99 | 1393 | Central | 287.41 | 0.07 | 0.06 | 285.56 | 0.08 | 0.54 |
| RW13 | 12 | 9 | OR | 45.33 | -121.66 | 1617 | Central | 333.09 | 0.05 | 0.04 | 472.20 | 0.05 | 0.63 |
| RW34 | 12 | 10 | BC | 50.97 | -122.80 | 1764 | Central | 373.83 | 0.05 | 0.04 | 199.83 | 0.05 | 0.62 |
| RW18 | 11 | 9 | OR | 45.18 | -117.30 | 2300 | Hybrid(E/C) | 411.69 | 0.07 | 0.05 | 984.59 | 0.07 | 0.52 |
| RW07 | 12 | 4 | BC | 51.25 | -117.47 | 1467 | Central | 418.80 | 0.06 | 0.04 | 267.35 | 0.06 | 0.58 |
| RW06 | 11 | 10 | BC | 51.76 | -119.93 | 1770 | Central | 423.00 | 0.04 | 0.03 | 190.31 | 0.05 | 0.69 |
| RW12 | 12 | 8 | BC | 49.61 | -115.48 | 1746 | Central | 457.84 | 0.04 | 0.03 | 342.23 | 0.04 | 0.69 |
| RW14 | 12 | 9 | OR | 44.04 | -121.74 | 1740 | Central | 476.99 | 0.06 | 0.04 | 554.64 | 0.06 | 0.64 |
| RW15 | 12 | 9 | OR | 42.90 | -122.07 | 1985 | Hybrid(W/C) | 612.56 | 0.13 | 0.12 | 4640.67 | 0.13 | 0.44 |
| RW16 | 11 | 9 | OR | 42.70 | -118.59 | 2587 | Central | 641.03 | 0.03 | 0.02 | 241.05 | 0.04 | 0.74 |
| RW08 | 12 | 8 | AB | 53.86 | -119.23 | 1862 | Central | 669.66 | 0.03 | 0.01 | 105.74 | 0.03 | 0.84 |
| RW26 | 12 | 10 | MT | 46.96 | -114.97 | 1726 | Eastern | 115.87 | 0.08 | 0.05 | 1784.79 | 0.08 | 0.62 |
| RW28 | 11 | 10 | MT | 47.47 | -112.93 | 1691 | Eastern | 184.00 | 0.08 | 0.05 | 1415.15 | 0.08 | 0.64 |
| RW19 | 12 | 10 | ID | 43.66 | -113.82 | 2524 | Eastern | 278.80 | 0.06 | 0.03 | 646.14 | 0.06 | 0.69 |
| RW17 | 11 | 10 | ID | 43.78 | -116.10 | 1826 | Eastern | 302.88 | 0.06 | 0.04 | 869.67 | 0.06 | 0.58 |
| RW29 | 11 | 10 | MT | 48.70 | -113.66 | 1839 | Eastern | 303.80 | 0.06 | 0.05 | 743.21 | 0.07 | 0.62 |
| RW27 | 12 | 8 | MT | 46.04 | -110.26 | 2187 | Eastern | 308.00 | 0.08 | 0.06 | 1061.16 | 0.08 | 0.55 |
| RW25 | 9 | 10 | WY | 43.71 | -110.88 | 2512 | Eastern | 396.88 | 0.07 | 0.05 | 750.56 | 0.07 | 0.53 |
| RW20 | 12 | 11 | NV | 41.84 | -115.43 | 2006 | Eastern | 503.44 | 0.05 | 0.03 | 591.59 | 0.05 | 0.74 |
| RW24 | 11 | 10 | WY | 42.74 | -109.21 | 2966 | Eastern | 586.10 | 0.06 | 0.05 | 725.76 | 0.07 | 0.58 |
| RW22 | 11 | 11 | UT | 40.58 | -111.62 | 2809 | Eastern | 677.94 | 0.06 | 0.04 | 1160.26 | 0.06 | 0.68 |
| RW21 | 11 | 10 | NV | 39.05 | -114.32 | 2541 | Eastern | 784.71 | 0.05 | 0.03 | 366.44 | 0.05 | 0.75 |
| RW23 | 12 | 10 | WY | 41.46 | -106.23 | 3012 | Eastern | 893.37 | 0.03 | 0.01 | 135.50 | 0.03 | 0.86 |

### Quantifying reproductive assurance

#### Greenhouse experiment

On January 16, 2023 we began a greenhouse experiment at the University of British Columbia by sowing 10–20 seeds from each maternal plant into 0.25 L square pots filled with moistened Sunshine mix #4 Professional Growing Mix (Sun Gro Horticulture, Agawam, MA). The 30 populations with field-collected seed (29 populations) or seed generated in the greenhouse via selfed hand–pollination from field collected cuttings grown to flowering in the greenhouse (1 population; see Supplemental Methods) were represented in our experiment, with 10–12 randomly chosen individual maternal lines per population. We completed two sowing cohorts (16 Jan & 6 Feb), sowing another five unique and randomly selected samples per population on the second sowing date. Seeds were germinated in pots on benches that were bottom-watered (flood benches) and top-misted with non-fertilized water three times/day for four weeks. Plants were grown in a controlled environment with photoperiod set to 15-hr days with compartment temperatures set to 12 ^◦^C min and 22 degrees ^◦^C max.

Pots with > 1 seedling were thinned 4–5 wk after sowing to retain the individual closest to the pot’s centre; a maximum of 11 pots per population were retained (*N* = 756 plants; Table 1). After thinning, we re-coded all samples so that population identity was unknown by data collectors. Once plants produced their first pair of true leaves, flood benches were switched to fertilizer-supplemented water (a custom blend containing micro- and macro-nutrients [N, P, K], designed to achieve a pH of 5.6-5.8). We constructed pollinator exclosures out of PVC pipe frames covered with bridal tulle (MEQUER brand). Plants were allowed to grow and mature in the absence of pollinator visitation or hand pollination. We marked three buds per plant beginning with the first bud and then every second flowering node. Buds were marked on the main stem of the plant by placing tape around the pedicel below the developing bud. Buds at each flowering node develop in sets of two and we haphazardly selected which to mark. Since floral ontogeny can vary during plant development and influence self-fertilization ability (Marshall et al., 2010), we tracked the order of the buds per plant to track floral ontogeny within focal plants. Once marked, we left buds to progress to anthesis and, after the flower matured and senesced (approx. 2 weeks), we carefully collected each individual fruit sample (calyx, carpels, pedicel) in a coin envelope taped closed on both ends to prevent seed loss. Note that mature fruits in *E. lewisii* dehisce within the upright, intact calyx and all seeds remain held within the fruit (carpels) and calyx so long as the plant is undisturbed, allowing us to effectively minimize potential seed loss in our experiment prior to sample collection.

One extreme temperature event (back-up power failure) occurred on May 27, 2023 where cooling was non-functional (though flood tables were still functional) and temperatures reached extreme levels for approximately 4 hours. However, at this stage of the experiment, all focal buds had been marked before the extreme temperature event and only eleven seed samples (out of a total of 765 samples across 255 plants) were not dehiscent before the extreme heat event.

After all seed samples were collected, we harvested the sexual reproductive (buds, flowers, fruits), asexual reproductive (rhizomes), and vegetative (above-ground tissue) biomass for each plant in separate paper bags, collecting counts of all sexual and asexual tissue types (i.e., number of buds, flowers, and fruits; number of rhizomes) before placing the separate tissue types in a drying oven set to 60 ^◦^C for 48 hours. Once dried, we weighed all of the biomass samples separately (Fisher Accu 4002d Precision Balance). We quantified clonal propagation by scoring probability of rhizome production and absolute count of rhizomes produced per plant. The experiment concluded on June 13, 2023.

#### Quantifying autonomous selfing ability

To quantify autonomous selfing ability, we counted any seeds that developed from marked buds in our experiment (i.e., seeds developed in the absence of pollinator visitation or hand pollination). We cleaned fruit samples of debris (i.e., calyx, carpel, pedicel), examined samples under a dissecting microscope, then—if seed was produced—counted seeds and weighed them (Mettler Toledo XPE105) after excluding aborted ovules or clearly inviable seeds.

#### Estimating range-wide reproductive assurance

We found a strong correlation between seed count and seed weight (mg) (*R*^2^ = 0.946) and therefore use seed count per fruit as a metric of selfing ability. We focus on rhizome count as our measure of clonal reproduction because we considered rhizome count to be a better metric of demographic success than rhizome weight. The seed and rhizome data sets were both zero-biased, with 61% and 59% of values being zero, respectively. We first analysed variation in seed production across the geographic range of *E. lewisii*, without the context of historical range expansion. Because a single zero-inflated model failed to converge, we analyzed two response variables in two models using the *lme4* (Bates et al., 2015) package in R, one for the probability of selfing (generalized linear mixed model with a binomial distribution) and one for seed number conditional on selfing (linear mixed effects model). Analogously, we next analysed the probability of producing rhizomes (generalized random effects model with a binomial distribution) and rhizome count (random effects model). In models of selfing ability, fruit ontogeny was coded as a fixed effect, whereas population and plant ID were coded as random effects. Since rhizome measurements came from a single plant (no repeated sampling), we did not include plant ID as a variable in models of clonal ability; only population was included as a random effect. To test for variation in reproductive assurance traits among populations across the geographic range, we performed a log-likelihood ratio test (*LRT*) using the *lmerTest* (Kuznetsova et al., 2017) package in R to assess whether the inclusion of the random effect “population” improved model fit.

### Phylogeography

#### ddRADseq genotyping

We extracted DNA from field-collected leaf tissue using a CTAB-chloroform protocol (full length protocol can be found at protocols.io) modified from Xin and Chen (2012). We added 100 ug/mL Proteinase K (NEB P8107S) to the CTAB Lysis buffer to reduce enzymatic oxidation and increased centrifugation speeds to 6000 rcf following DNA-CTAB complex precipitation. For DNA purification, we suspended 5 uL of MagAttract beads (Qiagen 1026901) in 55 µL of DNA plus TE Buffer (10 mM TRIS, 0.1 mM EDTA, pH 8.0) and performed multiple washes. 54 µL of the sample was transferred to a clean PCR plate. Purity was assessed using 2 µL of DNA elution on a Nanodrop 2000 (Thermo Scientific™), and DNA concentration was quantified with the Qubit 2.0 Broad Spectrum kit (Invitogen™) using 2 µL of DNA elution.

We then used a double-digest restriction-site associated DNA sequencing (ddRADseq) protocol to generate genome-wide sequence clusters (tags) for 372 individuals across 31 collection sites and 5 unique herbarium specimens, following the BestRAD library preparation protocol) and methods from Kolis et al. (2022). DNA samples were initially digested using restriction enzymes *PstI* and *BfaI* (New England Biolabs, Ipswich, MA; both methylation insensitive). After enzymatic digestion, sets of 48 individual DNA samples were labeled with unique in-line barcoded oligos, and each set pooled into a single tube. Pools were barcoded using NEBNext indexing oligos with a degenerate barcode, PCR amplified, and size selected (300-600 bp fragments) using a Bluepippin (Sage Science™) agarose gel. Libraries were prepared using NEBNext Ultra II library preparation kits for Illumina (New England Biolabs, Ipswich, MA). The completed libraries were paired-end sequenced (150 bp) on ^1^ lane of Illumina NovaSeqXPlus (Novogene Co., Ltd., Tianjin, China).

Illumina reads were demultiplexed and true PCR duplicates (same restriction site & same degenerate barcode) were removed using a custom Python script (Kolis et al., 2022). Adapters were trimmed and low-quality reads were removed using STACKS function `process shortreads` (Catchen et al., 2013, 2011). Reads were mapped to the *E. erubescens* version2 reference genome (http://mimubase.org/FTP/Genomes/LF10g_v2.0/) using BWA MEM (Li and Durbin, 2009) and indexed with SAMtools (Li et al., 2009). SNPs were called using GATK (McKenna et al., 2010) for aligned sequence data. After alignment and quality filtering (quality> 30, maximum missing data = 0.2), we retained 186, 746 variable sites. We filtered for sites with mean coverage g10 & f55× and individuals with mean depth g10×, retaining 120, 681 variable sites and 356 individuals representing all 31 collection sites (9–12 individuals per site; Table 1) and three out of five Alaskan herbarium samples (which were then pooled to be considered as one population) in our genomic dataset. For analyses of population structure we filtered minor allele frequency (*ma f*) > 0.1 with bcftools (Danecek et al., 2021) and linkage disequilibrium with PLINK (Purcell et al., 2007) and retained a total of 13, 055 variable sites. For analyses of genetic diversity, we used genomic data where we did not filter for minor allele frequencies or linkage disequilibrium.

#### Spatial population genetic structure

To analyze population genetic structure we generated a dataset for principal component analysis (PCA) using the `pca` function in PLINK (Purcell et al., 2007) to summarize the overall genetic similarity of individuals and ran the program ADMIXTURE (Alexander et al., 2009) for *K* = 1–30 to generate individual-level ancestry assignments.

We used the software “Fast Effective Estimation of Migration Surfaces” (FEEMS) (Marcus et al., 2021; Petkova et al., 2016) to examine spatial patterns of gene flow across the species geographic range by estimating migration rates between populations that deviate from expectations under isolation-by-distance. This method visualizes regions of the landscape that may facilitate or hinder gene flow, allowing us to contextualize emergent patterns of population genetic structure. To optimize our FEEMS output, we performed leave-one-out cross-validation over a grid of *λ* values (1*e*^−6^ – 1*e*^2^; model tuning parameter) to select the model fit with the lowest cross-validation error.

#### Genetic differentiation

We calculated pairwise genetic distances (Weir and Cockerham’s *F_ST_*) between all populations and ancestral clusters using functions from the R package *hierfstat* (Goudet and Jombart, 2022) within the R package *graph4lg* (Savary et al., 2021). For all analyses involving comparisons within and between ancestral clusters, we removed putative hybrid populations (two in total) to prevent obscuring signals of relatedness.

We generated topographic distances between sampled populations using the open source website OpenTopography and R package *elevatr* (Hollister et al., 2023) to generate a topographic raster layer of the area of the entire geographic range of *E. lewisii*. We then used the R package *topoDistance* (Wang, 2020) to calculate a pairwise matrix of topographic distances between all sampled populations. To test for patterns of isolation-by-distance within distinct spatial population genetic clusters, we used the R packages *ade4* (Dray and Dufour, 2007) and *vegan* (Oksanen et al., 2022) to run separate Mantel tests per ancestral cluster, with 9999 permutations each.

#### Genetic and Nucleotide diversity

We used the R package *snpR* (Hemstrom and Jones, 2023) to calculate the following measures of genetic diversity within each distinct population: mean *H_o_*, mean *H_e_*, and count of private alleles. We used PLINK to calculate individual-level inbreeding coefficients (*F_IS_*) to test the relationship between our measure of population-level selfing ability in the greenhouse with genetic estimates of inbreeding in nature.

To investigate nucleotide diversity, we generated an “all sites” vcf (including both variant and invariant sites) using GATK (McKenna et al., 2010) and filtered for depth and missing data in the same way as above (retained 189,823 sites) to avoid biasing our estimates of nucleotide diversity (Korunes and Samuk, 2021). We then used the software *pixy* to calculate average nucleotide diversity within (*π*) and between (*d_xy_*) populations and ancestral clusters over a 1 Mb window per chromosome. We then averaged the per-region values to generate a per-individual value and finally generated a population mean.

#### Identifying putative origins of range expansion

We used the directionality index, *ψ*, and the time distance of arrival (TDoA) method (Peter and Slatkin, 2013) to infer the most likely locations of the origin of range expansion within unique population genetic clusters. The directionality index statistic, *ψ*, is based on range expansion theory involving serial founder effects (Slatkin and Excoffier, 2012) and uses asymmetries in frequencies of shared derived alleles (excluding alleles that are not shared among populations because they have either been lost or have reached fixation in a given population) to detect the direction of colonization (donor vs. recipient) between two populations of interest. Specifically, populations further from the origin are expected to have undergone more founder events (sampling effects) and experienced more sampling drift, so shared derived alleles are expected to be at relatively higher frequencies in the more recently colonized recipient population compared to the older donor population. To calculate *ψ*, we phased the genomic data with the reference genome (sister species *Erythranthe erubesens*) as the ancestral allele and calculated the directionality index and *ψ* statistic for all pairwise combinations of populations using the `calc directionality()` function in *snpR*.

To infer the most likely locations of the origin of range expansion within unique population genetic clusters, we combined the directionality index results with the sampling coordinates (Table 1) for each population in the TDoA method (Peter and Slatkin, 2013). The TDoA approach identifies the origin(s) of the expansion by locating the deme(s) that explain the highest proportion of variation in the correlation between pairwise *ψ* differences and pairwise geographic distances differences (He et al., 2017; Peter and Slatkin, 2013) by performing regression analyses between these sets of pairwise differences (largest *R*^2^-values are identified as geographic location closest to the potential origin). We implemented the TDoA method using archived R scripts from Koski et al. (2019). To further explore the assignment of putative refugia in the context of historical range expansion (i.e., serial founder events Austerlitz et al. 1997; Hewitt 2000) vs. contemporary ecology (i.e., the ’centre–periphery hypothesis’ or the ‘abundant centre hypothesis’ Pironon et al. 2017) influencing spatial patterns of genetic diversity, we tested how the relationship between genetic and nucleotide diversity changed on average with latitude, using linear and quadratic terms in our models and selecting the model with lowest AIC score. Support for a linear fit would match a unidirectional expansion from southern refugia, whereas support for a quadratic fit would match the prediction for the abundant centre hypothesis.

#### Testing reproductive assurance hypotheses

We used a zero-inflated model to test the relationship between autonomous selfing ability and distance from the putative origin of range expansion, controlling for ancestral cluster and repeated sampling of focal plants as random effects. We also tested the relationship between clonal propagation (measured as probability of rhizome production and rhizome count) and distance from putative origins of range expansion. For rhizome data, since a zero-inflated model failed to converge, we analyzed two response variables in two models, one for the probability of rhizome production (generalized linear mixed effects model with a binomial distribution) and one for rhizome number conditional on rhizome production (linear mixed effects model), controlling for ancestral cluster. For the model of probability of producing rhizomes, we scaled the predictor variable (distance from origin) to achieve model convergence.

### Analysis of reproductive investment

To test if individuals varied in their investment in floral display, we calculated the ratio between floral reproductive and aboveground vegetative biomass. We ran a linear mixed effects model to test whether investment in floral reproduction varied significantly with distance from putative refugia, with the prediction that if relative investment in floral biomass decreased on average with distance from the origin of expansion that this would be consistent with support for an increasingly “selfing-syndrome”-like phenotype (Goodwillie et al., 2010) towards the range edge.

## Results

### Geographic variation in autonomous selfing and clonal growth

Across space and without the context of historical range expansion, we found all models that included population as a random effect performed better (i.e., explained more of the variance in the associated response variable) than models without population included as a random effect Fig. 2 (probability producing selfed seed: *LRT* = 88.01, *p* < 0.0001; seed count per selfed fruit: *LRT* = 4.02, *p* = 0.04; probability producing rhizomes: *LRT* = 12.42, *p* < 0.0001; rhizome count: *LRT* = 10.46, *p* = 0.001). There were no effects of ontogeny of flowers on variation in probability of selfing (*χ*^2^ = 2.19, *d f* = 2, *p* = 0.33) or absolute seed count (*χ*^2^ = 1.04, *d f* = 2, *p* = 0.6). Finally, we found evidence that greenhouse measures of selfing ability reflected patterns of inbreeding in the field, where population-level probability of producing selfed seed explained 16% (*R*^2^ = 0.16) of the variance in population-level inbreeding coefficients (*β* = 0.21, SE =, 0.08, *p* = 0.015) (Fig. S2). Across all samples, inbreeding coefficients were high (mean: 0.70 ± 0.009) (Table 1).

**Figure 2:**
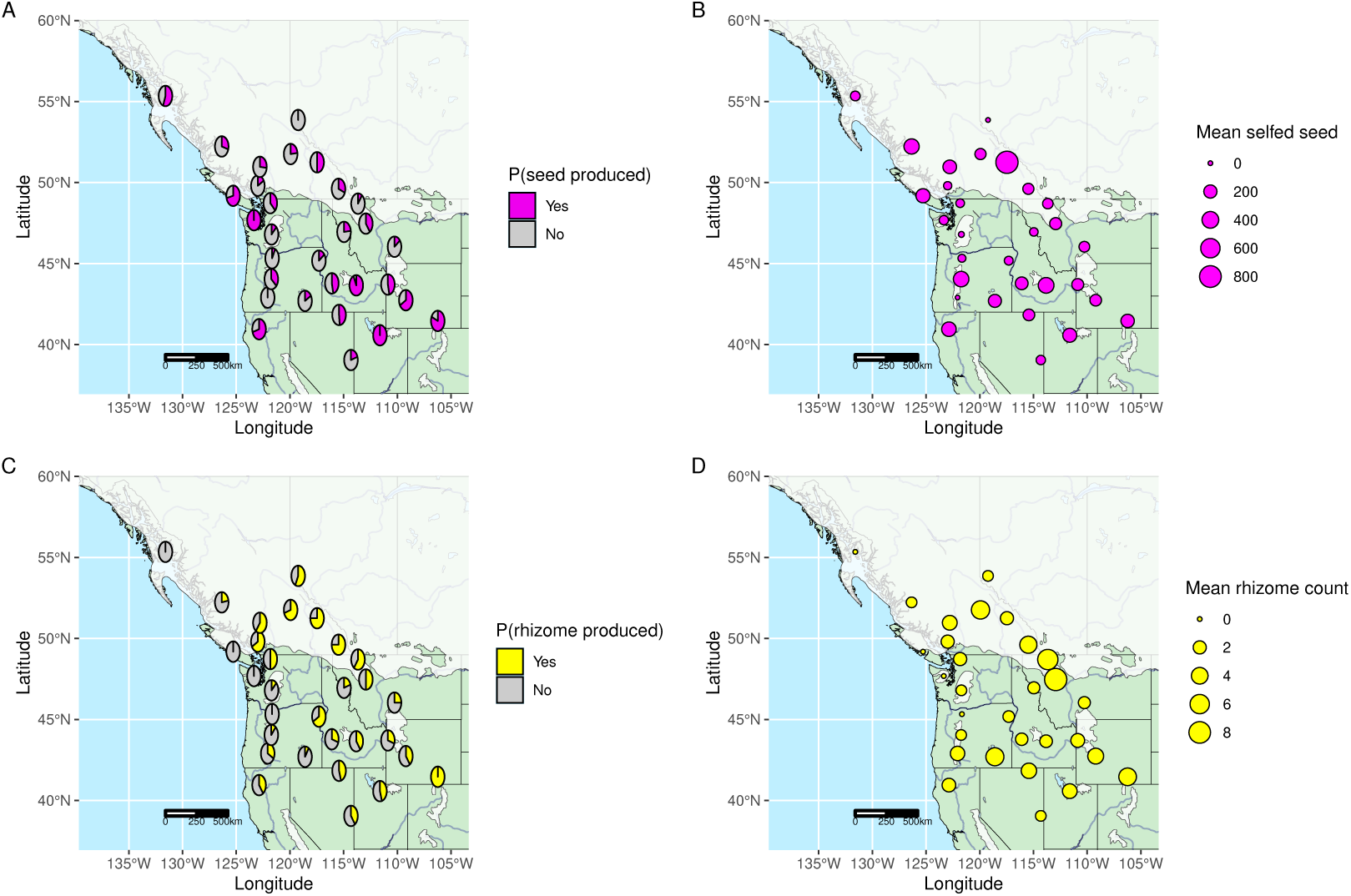
Greenhouse measure of reproductive assurance in *E. lewisii* exhibits range wide variation. Range wide maps with last glacial maxima of the Cordilleran ice sheet (white overlay) (Dalton et al., 2020) depicted where *E. lewisii* seed was sampled across 30 populations used in our greenhouse survey of reproductive assurance. A) Percent population-level probability of selfing, B) Absolute mean number of selfed-seed produced (i.e., values were calculated without zeros included); populations with a mean selfed-seed count of zero indicate populations with no detected selfing ability in the greenhouse. C) Population-level probability of producing rhizomes, and D) Absolute mean number of rhizomes produced (i.e., values were calculated without zeros included); populations with a mean rhizome count of zero indicate populations with no detected rhizome production in the greenhouse. For panels B) and D), size of point reflects number of seeds or rhizomes produced. For further detail on variance in absolute seed and rhizome counts produced, see Fig. S3.

### Spatial genetic structure

In the PCA, PC1 explained 27.4% and PC2 explained 19.8% of variation in population genetic diversity, with visually distinct separation of three main clusters along PC1 and PC2 and two potentially admixed clusters (Fig. 3A). Based on the visual representation of genetic diversity in the PC plot, we investigated ancestry assignments in ADMIXTURE analysis with a cluster value of *K* = 3 (Fig. 3B). We also examined the cross-validation error plot from the ADMIXTURE analysis, however; clustering algorithms can overfit data easily, tending to overestimate the numbers of discrete clusters present, and caution is recommended when interpreting these inferred values of *K* (Pritchard et al., 2000) (in our case, *K* = 17, 15, & 16; see Fig. S4). For these reasons, we based our ancestral cluster assignment on the visual structure in PC space. With *K* = 3, our ADMIXTURE analysis identified two populations assigned hybrid ancestry (RW15: 42.9047, −122.0702 & RW18: 45.1817, −117.3038; Table 1) that discretely corresponded with the two intermediate clusters along PC1 and PC2 in our PCA plot (Fig. 3A) and therefore we excluded these two populations from further analyses of discrete ancestral clusters.

**Figure 3:**
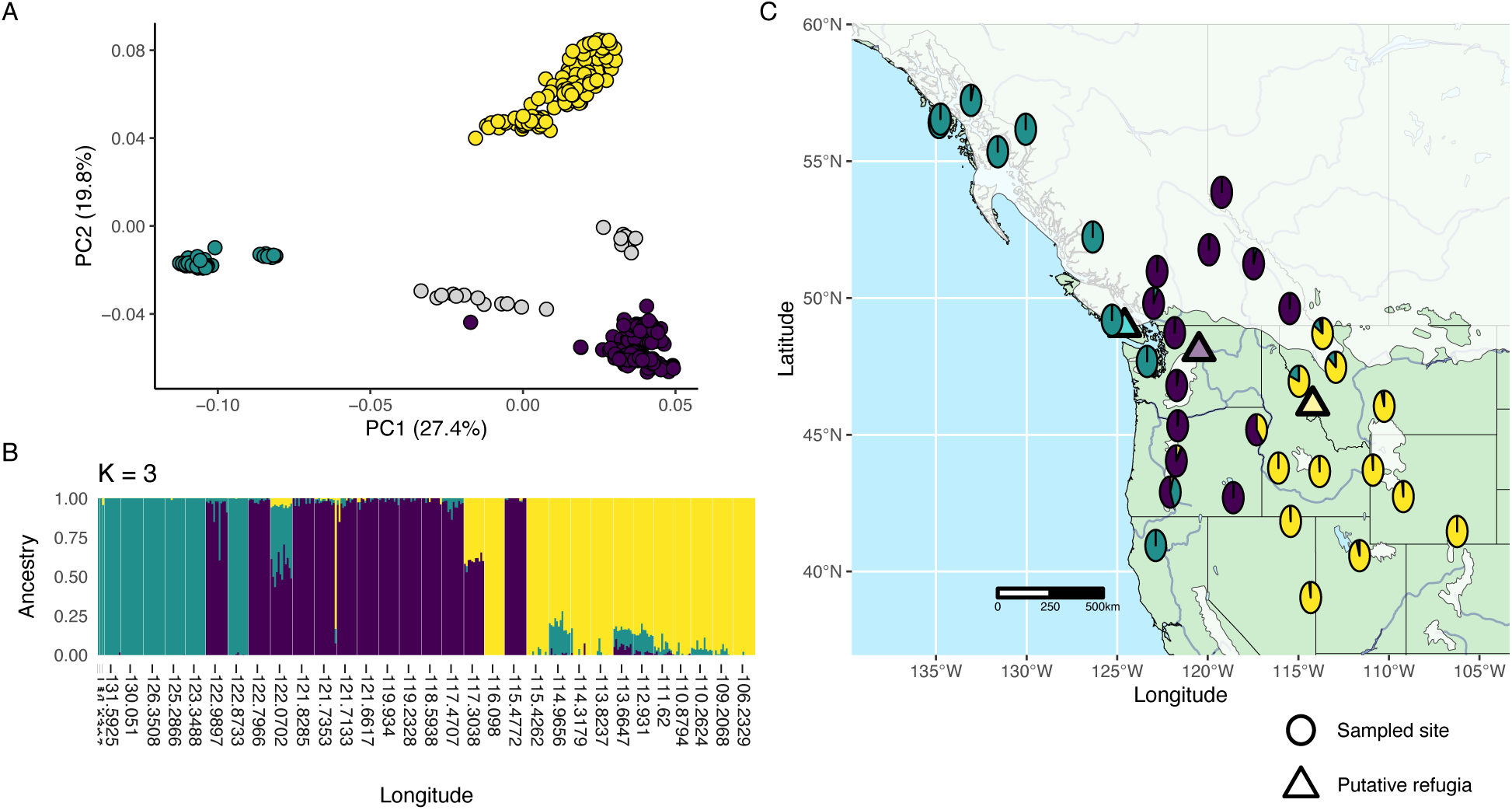
Distinct spatial population genetic structure detected across longitude. A) PCA plot of first two PC axes (PC1 explains 27.4% and PC2 explains 19.8% of variation) summarizing genetic structure/overall genetic similarity of individuals and depicting three distinct genetic clusters and two putative hybrid clusters (points coloured by majority ancestry assignment from panel B) where grey points represent putative hybrid populations [lon: −122.0702; lon: −117.3038]). B) ADMIXTURE plot with *K* = 3, populations ordered by longitude. C) Mean population-level ancestry proportions (from B) plotted on a map encompassing the entire geographic range of *E. lewisii* with last glacial maxima of the Cordilleran ice sheet (white overlay; Dalton et al. 2020) and locations of three putative origins of range expansion (from TDoA analysis) for each genetic cluster (coloured triangles correspond to ancestral cluster colours from panel A and B).

We then mapped the average ancestry assignments per population in geographic space and resolved a spatial pattern among cluster assignments that mapped with longitude (Fig. 3C). We calculated genetic differentiation and nucleotide diversity within and divergence among ancestral clusters and found strong divergence between the Western and Central clusters (*F_ST_* = 0.67; *d_xy_* = 0.25), Western and Eastern (*F_ST_* = 0.62; *d_xy_* = 0.26) clusters, and moderate divergence between the Central and Eastern clusters (*F_ST_* = 0.25; *d_xy_* = 0.11). All ancestral clusters had similar levels of nucleotide diversity (Western: *π* = 0.102; Central: *π* = 0.072; Eastern: *π* = 0.095). Within each genetic cluster, we detected significant patterns of isolation-by-distance, a pattern that can emerge from sequential founder events and limited migration during range expansion (Western: *r* = 0.66, *p* < 0.01; Central: *r* = 0.71, *p* < 0.0001; Eastern: *r* = 0.63, *p* < 0.0001) (Fig. S5).

Our investigation of the estimated effective migration surface suggested uneven migration across the landscape (Fig. S6) concordant with patterns of population structure (Fig. 3C). Areas of low migration/gene flow between clusters corresponded with landscape features that likely limit dispersal such as the Coast/Cascade mountains between the Western and Central clusters and areas within the Rocky Mountain range between the Central and Eastern clusters.

### Inference of historical origins of range expansion

We subset pairwise contrasts of the metric *ψ* by ancestral cluster and estimated three refugia (one per cluster) for the origin of range expansion using the TDoA method (3C) located in Western Vancouver Island, BC (Western cluster: −125.2925, 49.3589), North Cascades, WA (Central cluster: −120.4843, 48.0656), and Bitterroot mountains, MT (Eastern cluster: −114.206, 46.0813) (Fig. S7). From the TDoA analysis, we detected significant relationships between pairwise differences in the directionality index *ψ* and geographic distances between populations for the Central (*R*^2^ = 0.62, *p* < 0.0001) and Eastern (*R*^2^ = 0.42, *p* < 0.0001) clusters, but did not detect a significant relationship within the Western cluster (*R*^2^ = 0.25, *p* > 0.05), suggesting significant signals of range expansion from the putative origin in two out of three clusters. Overall, consistent with the prediction of the abundant centre hypothesis, we found that quadratic models generally fit the data better than linear models (i.e., lower AIC values, higher *R*^2^ values) for the relationship between measures of genetic diversity and latitude, suggesting genetic diversity was highest on average in the centre of the geographic distribution (Table S1, Fig. S8; but see nucleotide diversity with almost identical AIC values).

### Relationship between spatial patterns in reproductive assurance and historical pathways of range expansion

To test our main hypothesis of the effect of historical range expansion on spatial patterns of reproductive assurance, we examined how autonomous selfing and rhizome production varied with distance from putative refugia. A zero-inflated model showed the probability of producing selfed seed increased with increasing distance from inferred refugia, by an average magnitude of ∼0.25 at the maximum distance compared to the inferred origin (*β* = 0.0014, SE = 0.0004, *p* < 0.001; note that the sign of coefficient is reversed from default outputs to express the probability of making selfed seed rather than probability of failing to make selfed seed) (Fig. 4A). We did not find support for the hypothesis that variation in the absolute number of selfed seed produced varied with distance from refugia, as demonstrated by the conditional portion of our model (*β* = 0.0002, SE = 0.0004, *p* = 0.511) (Fig. 4B). A generalized mixed effects model showed showed the probability of producing rhizomes increased with increasing distance from the inferred refugia by an average magnitude of ∼0.3 at the maximum distance compared to the inferred origin (unscaled coefficient: *β* = 0.0015, SE = 0.0006, *p* = 0.013) (scaled coefficient: *β* = 0.352, SE = 0.142, *p* = 0.013) (Fig. 4C). We did not detect a significant relationship between distance from inferred refugia and rhizome count per individual (*β* = −0.001, SE = 0.001, *p* = 0.41; Fig. 4D).

**Figure 4:**
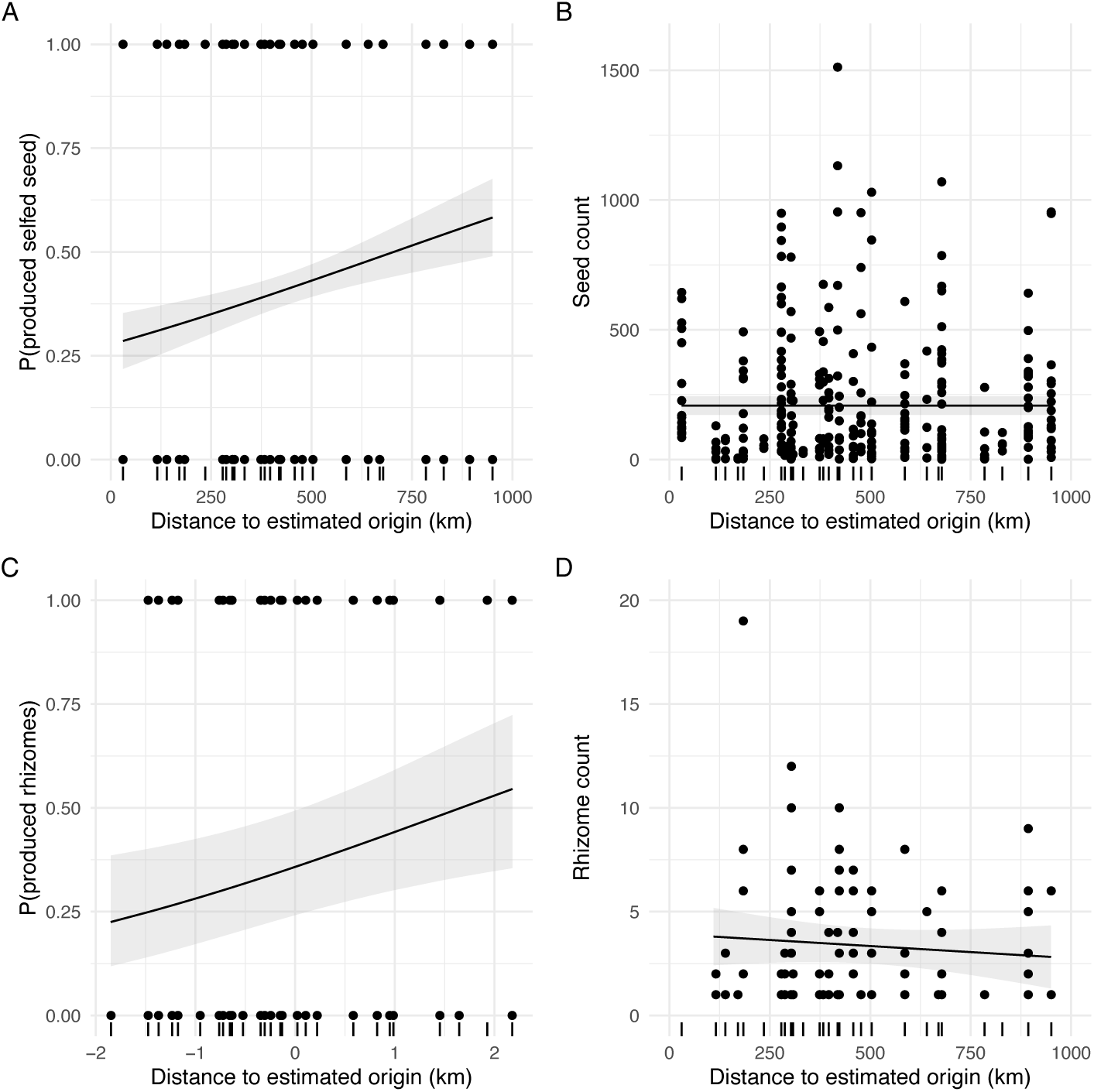
The probability of both sexual and asexual reproductive assurance increases on average with increasing distance from inferred refugia/origin of expansion. A) Average model output from zero-inflated portion of model (with sign of estimate reversed) with per-sample binomial response data plotted showing a positive relationship between probability of selfing and distance from putative refugia. B) Average model output from conditional portion of zero-inflated model (values > 0) with per-sample seed count data plotted showing no relationship between number of autonomously produced seed and distance from putative refugia. C) Average model output from generalized linear mixed effects model with per-plant binomial response data plotted showing a positive relationship between probability of producing rhizomes and distance from putative refugia. Note, the values on the x-axis were scaled for this model. D) Average model output from linear mixed effects model with per-plant rhizome count data (values > 0) plotted showing no relationship between number of rhizomes produced and distance from putative refugia. All regression lines represent average model fit, controlling for ancestral cluster (A, B, C, D) and repeated measures within plants (A, B), with 95% confidence intervals depicted via gray shading. Points in panels A and C represent probabilities of individual samples producing selfed seed (A) or rhizomes (C) where “0” indicates never producing selfed seed or rhizomes and “1” indicates production of seed or rhizomes. Dashes on the x-axis represent locations of sampled populations.

In support of our greenhouse-based measures of reproductive assurance via uniparental reproduction, our genetic measure of population-level inbreeding *F_IS_* significantly increased on average with increasing distance from putative refugia (*β* = 1.861*e*^−04^, SE = 5.136*e*^−05^, *p* < 0.01). We did not detect any differences in relative investment in floral vs. vegetative growth with distance from the putative origin (*β* = 7.98*e*^−05^, SE = 4.44*e*^−05^, *p* = 0.073) (Fig. S9).

## Discussion

Using Northern pink monkeyflower as a focal study system, we documented substantial spatial variation in reproductive assurance via both self-pollination and clonal growth, identified three putative independent refugia from which populations recolonized formerly glaciated areas, and related probability of reproductive assurance to pathways of post-glacial recolonization. Consistent with the hypothesis that reproductive assurance evolves during range expansion, we found that the probability of producing selfed seed and the probability of producing clonal rhizomes significantly increased on average with distance from the putative refugial origin of range expansion. This follows hypotheses underlying Baker’s law, where demography during colonization selects for an enrichment in the capacity for uniparental reproduction in colonizing species under mate and/or pollinator limitation (Pannell et al., 2015).

While previous work has tested the evolution of reproductive assurance in peripheral populations (Busch, 2005; Herlihy and Eckert, 2005; Kennedy and Elle, 2008), the present study contributes to a growing literature of empirical tests demonstrating evidence for the role of historical range expansion shaping plant mating system evolution in autonomous selfing ability (Koski et al., 2019; Willi et al., 2018; Zeitler et al., 2023) and is the first to our knowledge to test these specific effects on clonal reproductive assurance. Our focal species, *E. lewisii*, differs in sexual reproduction from those in previous studies in that it maintains a relatively large, showy floral display range-wide (compared to Willi et al. 2018 [*Arabidopsis lyrata*] & Zeitler et al. 2023 [*Arabis alpina*]) and has no well-known active mechanisms for delayed selfing (unlike protandry [i.e, earlier maturation of male sexual organs] in *Campanula americana* [Koski et al. 2019]). Given the growing evidence, we suggest post-glacial range expansion may be a general mechanism shaping geographic distributions in mating system evolution for numerous flowering plant species encompassing variation in floral display size, mating system traits, and life history strategies. Below, we also expand on alternative hypotheses that could contribute to our detected relationship between pathways of historical range expansion and reproductive assurance.

### Evolution of reproductive assurance following pathways of historical range expansion

*Erythranthe lewisii* has the capacity for both sexual and asexual uniparental reproduction (i.e., autonomous selfing and clonal propagation) and our results match the pattern described by Baker (1955) where species’ capacity for self-fertilization (and/or purely vegetative propagation) is predominant among colonizing individuals, increasing towards range edges while concurrently maintaining the capacity for outcrossing (i.e., no known self-incompatibility mechanisms) (Pannell et al., 2015). The patterns we detected suggest that selection for both selfing and clonal propagation during colonization occurred during range expansion. Alternatively, had we detected changes in only one component, or concomitant decreases in one while the other increased, this would have suggested differences in resource allocation (i.e., different relative investment in sexual vs. vegetative tissues) or a trade-off.

We detected range-wide variation in all greenhouse-measured reproductive assurance traits, but only the probabilities of producing any selfed-seed and rhizomes were related to inferred range expansion from historical refugia; numbers of selfed seeds or rhizomes, when they could be produced, varied among populations but did not change linearly with distance from refugia. Future work can examine range wide differences in ovule number and pollen production/viability, and how these vary in the context of historical range expansion to influence absolute number of seeds produced via selfing. As for Baker’s Law, the original paper stated a single propagule from a self-compatible individual is sufficient to start a sexually-reproducing colony (Baker, 1955), suggesting probability of autonomous reproduction is the most important metric for colonization success.

Overall, our greenhouse experiment suggests selection for autonomous selfing and clonal growth occurred during historical range expansion, though the effect of current ecological conditions warrants further investigation. Future work should quantify the role of pollen limitation in natural populations (Busch, 2005) by conducting pollinator surveys and experimental crosses in the field to evaluate the relative role historical range expansion has played in shaping geographic patterns in mating system diversity. Alternative explanations for elevated clonality with pathways of range expansion related to current ecological conditions can also include pollinator abundance (Vallejo-Maŕın et al., 2010) and seasonality (Villegas, 2001).

### Origins of expansion

Inferred ancestral genetic clusters were well supported in terms of genetic differentiation and geographic barriers to dispersal (Fig. S5, Fig. S6). Low levels of nucleotide diversity within clusters are likely related to generally low levels of expected heterozygosity and high levels of inbreeding across the range of *E. lewisii* (Table 1).

We found statistical support for signatures of range expansion from a putative origin for two out of three ancestral clusters, all of which were located on habitat that was ice-free during the last glacial maxima (∼20,000 years ago) (Dalton et al., 2020). Signatures of range expansion were not statistically supported for the Western cluster and this could be due to relatively smaller sample size within this cluster (Table 1) and more isolated habitat (Fig. S6) potentially leading to reduced power to detect genetic signatures of expansion.

Overall, we found models with a quadratic term included were better at explaining the relationship between levels of genetic diversity across the latitudinal range of *E. lewisii* than linear models (Table S1). This demonstrates that, on average, levels of genetic and nucleotide diversity were greatest at mid-latitudes, a pattern that is superficially consistent with the abundant-centre hypothesis, a contemporary ecological explanation for spatial patterns in genetic diversity that states that genetic variation of a species should decrease from the centre to the edge of its geographic range (Brown, 1984) . However, identified refugia were located at mid latitudes, not the southern portions of the species range. Historical refugia should have the highest levels of genetic diversity based on age, with serial founder events causing losses of ancestral diversity as new populations are founded during post-glacial range expansion (Austerlitz et al., 1997; Hewitt, 2000). Thus, the mid-latitude peak in genetic diversity cannot discriminate between the abundant centre hypothesis and historical range expansion from interior (rather than southern) refugia.

However, in contrast to contemporary ecological explanations, further support for the historical processes underlying the identification of three glacial refugia existing at mid-latitudes comes from growing evidence for middle latitude origins in North America, Europe, and Asia (Bai et al., 2010; McLachlan et al., 2005; Peterson and Graves, 2016; Petit et al., 2003; Prior and Busch, 2021) and a meta-analysis of the locations of historical glacial refugia in Northwestern North America (Shafer et al., 2010). This follows what we know about the unique and complex glacial history and physical geography (i.e., multiple mountain ranges) of the region, including the existence of multiple refugia for a given species, that challenges the simpler view of consistent single southern origins of post-glacial expansion (Petit et al., 2003; Shafer et al., 2010). The aforementioned meta-analysis demonstrated there is consensus among previous studies that historical refugia for multiple taxa exist in all three regions where our putative refugia were located (support for major refugia in Vancouver Island, BC and Rocky Mountains/Bitterroot mountains, MT; support for cryptic refugia in Cascade Mountains, WA). To complement what we know from comparative phylogeographic studies, additional independent lines of evidence for historical environmental suitability across the species’ range using pollen data and hindcast niche modelling (Waltari et al., 2007) could further evaluate the assignment of refugia.

### High levels of inbreeding range-wide

Increased selfing can decrease the effective population size and the amount of standing genetic variation within a population (Barrett and Harder, 2017), which can have negative consequences for long-term population persistence (Goldberg et al., 2010). Across the range, we found that expected heterozygosity was consistently higher than observed heterozygosity and values for population-level inbreeding coefficients were generally high (Table 1), both indicative of nonrandom mating and low effective population sizes. *Erythranthe lewisii* possesses a large showy floral display, commonly associated with predominantly outcrossing plant species (Goodwillie et al., 2010), however our genetic results suggest a high degree of non-random mating within populations that could be attributed to self-fertilization (Igic and Kohn, 2006), the latter which can be increased by clonal propagation (geitonogamy via clonal genets). Future studies could discern the relative contributions of autonomous vs. geitonogamous selfing on inbreeding levels in nature by conducting floral manipulations in the field (e.g., manipulating individual flowers to only allow autonomous selfing vs. forced outcrossing [anther removal]) combined with genetic markers to estimate differences in seed production compared to controls (Eckert, 2000).

While high levels of inbreeding can lead to reduced fitness due to inbreeding depression (Charlesworth, 2006), over time low levels of heterozygosity due to inbreeding may reflect the purging of deleterious recessive mutations from populations (Byers and Waller, 1999), ultimately resulting in populations overcoming the negative effects of inbreeding depression (Crnokrak and Barrett, 2002; Pujol et al., 2009). Additionally, drift load can accumulate during range expansion (Peischl et al., 2013; Peischl and Excoffier, 2015), and the tension between accumulated drift load and purged inbreeding depression on fitness can be examined by conducting within- and between-population experimental crosses and examining heterosis in the context of historical range expansion (Koski et al., 2019; Willi et al., 2018; Zeitler et al., 2023). Individuals that have been able to purge deleterious mutations via sexual reproduction (including via geitonogamous selfing) would be at an advantage to maintain those genotypes via clonal reproduction, perhaps explaining the elevated probability for both forms of reproductive assurance. Purging could also predispose individuals to express higher levels of autogamy without negative genetic consequences. Future studies examining the extent of inbreeding depression vs. drift load among populations in the field can examine how these spatial patterns reflect pathways of historical range expansion (Koski et al., 2019).

### Conclusions

Using a combination of range-wide field sampling, a large-scale greenhouse experiment, and phylogeographic reconstruction, we present strong evidence for the role of historical range expansion driving contemporary spatial patterns in both sexual and asexual reproductive assurance in *E. lewisii*. Our study joins a handful of others that use phylogeographic reconstructions and greenhouse-based estimates of reproductive assurance (Koski et al., 2019) or *a priori* knowledge of spatial variation in population-level mating system (Willi et al., 2018; Zeitler et al., 2023) to conduct high-caliber tests of Baker’s law (Baker, 1955). Our results demonstrate relatively low heterozygosity and high inbreeding coefficients range-wide, and this could indicate high levels of clonal propagation and potential interactions between the evolution of selfing and purging of deleterious mutations that can be examined in the future via spatial patterns of inbreeding depression and drift load. For a complete picture of the drivers of spatial patterns in reproductive assurance in *E. lewisiii*, future studies should also investigate the roles of pollinator and mate limitation in the field. In light of our relatively recent ability to successfully collect the data necessary to directly test these questions in ecology and evolution research, the growing evidence in the literature suggests reproductive assurance during post-glacial range expansion may be a common mechanism maintaining diversity in plant mating systems; future investigations across a greater diversity of plant species will further inform our understanding of its generality.

## Author contributions

MUC and AA conceived of the study and designed the field sampling. MUC and KT conducted fieldwork. MUC conducted all greenhouse work, lab work, and generated and analyzed all data. MUC wrote the first draft of the manuscript with input and contributions from all authors. Funding from NSERC DG to ALA (RGPIN-2022-03113). Research grant funding was provided to MUC from The American Philosophical Society (Lewis and Clark Fund for Exploration and Field Research), The American Society of Naturalists (Student Research Award), and The Botanical Society of America (Graduate Student Research Award).

## Acknowledgements

Thanks to Michael Whitlock, Jennifer Williams, and Loren Rieseberg for helpful comments on an earlier draft of this manuscript. Thanks to Francisco Henao-Díaz, Julian Heavyside, Olivia Rahn, Graydon Gilles, Xianyu Yang, Samantha Brown, Bill Baccus, Graeme Keais, and Trevor Goward for assistance with fieldwork. Thanks to Jill Jankowski for providing critical field equipment. Thanks to many different private and public land managers, employees, and botanical experts including Joni Johnson (Tongass National Forest), Justin Fulkerson & Matthew Carlson from The Alaska Center for Conservation Science, Gretchen Baker (Great Basin NP), Caleb Catto (@calebcatto on iNat), Harvey Thomassen, Carol Thomassen, & Lise Baille, and Trevor Goward & Curtis Bjork for assistance (please see “Supplementary Acknowledgements” for complete fieldwork acknowledgments). Thanks to Jacob Arogones and Lauren McBurnie for assistance in the greenhouse and to Melina Biron and UBC Plant Care Services for support in the greenhouse. Thanks to Linda Jennings and Spencer Goyette at the UBC Herbarium and Stefanie Ickert-Bond at the University of Alaska Herbarium for providing leaf tissue samples for DNA extraction. Thanks to the Rieseberg and Whitton Labs at UBC for space and equipment to complete DNA extractions. Thanks to Dylan Moxely at UBC for support with DNA extractions and Colette Berg, Lila Fishman, Tim Wheeler, David Xing, and the University of Montana Genomics Core for providing reagents, equipment, and support to complete ddRADseq library preparation. Thanks to the Williams lab for space and equiptment to quantify autonomous seed set. Thanks to Julia Kreiner and Fernando Hernández for assistance with genomic analyses. This research was enabled in part by support provided by the Digital Research Alliance of Canada (https://alliancecan.ca/en).

## Data accessibility

All data and analysis code used in this article will be deposited in Zenodo upon publication. Sequencing data will be deposited in the NCBI Sequence Read Archive.

## Supplementary Methods

### Greenhouse experiment

Note, for the site sampled near Grand Cache, AB (Fig. 1), we sampled leaf tissue and collected live stem cuttings in the field in late July 2021, propagated cuttings to flowering in the green-house (by coating stems in rooting hormone [ProMix Stim-Root Rooting Powder] and transplanting stems into 0.25 L square pots filled with wetted-down Sunshine mix #4 Professional Growing Mix (Sun Gro Horticulture, Agawam, MA) and bottom watering three times/day with fertilized water in a controlled environment with photoperiod set to 15-hour days with compartment temperatures set to 12 ^◦^C min and 22 degrees ^◦^C max.), and performed within-flower crosses to generate seed samples (i.e., this is the only population in which seed was not directly collected from the field).

## Supplementary acknowledgments

We would like to thank the following additional people and organizations for assistance in permitting fieldwork: Karen Dillman (Tongass National Forest), Kitty LaBounty (University of Alaska Southeast), Andrea Zemenak (Wells Gray Provincial Park), Geoff Popowich & David Whiteside (Garibaldi Provincial Park), Ben Grove (Wells Gray Provincial Park), Matthew Dubeau (Olympic National Park), Tara Chestnut (Mount Rainier), Karin Hawley (Mt. Hood), Heather Jackson (Three Sisters Wilderness - Deschutes National Forest), Melynda Beam (Three Sisters Wilderness - Deschutes National Forest), Jen Hooke (Crater Lake), Wendy Wayne (Crater Lake), Kyle Wanner (BLM: Steens Mountain Wilderness), Joshua Nicholes (Humboldt-Toiyabe National Forest), Wendy Markham (Humboldt-Toiyabe National Forest), Weston McPhie (Uinta-Wasatch-Cache National Forest), Marti Aitken (Medicine Bow-Routt National Forest), Christie Schneider (Medicine Bow-Routt National Forest), Tara Carolin (Glacier National Park), Natalie Stafl (Glacier National Park Canada), and Lusetta Sims (Klamath National Forest).

Fieldwork was conducted under the following permits: Olympic National Park (NP) (OLYM-2020-SCI-0043), Mount Rainier NP (MORA-2021-SCI-0009), Glacier NP (GLAC-2020-SCI-0020), Glacier NP Canada (GLA-2020-35880), Crater Lake NP (CRLA-2020-SCI-0002), Great Basin NP (GRBA-2021-SCI-0007), Grand Teton NP (GRTE-2021-SCI-0048), Helena-Lewis & Clark Nation Forest (NF) (024968), Tongass NF (003790), Uinta-Wasatch-Cache NF (014794), and Medicine Bow-Routt NF (027183). Letters of Permission or Authorization granted from: Boise NF, Salmon-Challis NF, Humboldt-Toiyabe NF, Wallowa-Whitman NF, Mt. Hood NF, Steens Mountain Wilderness BLM, Three Sisters Wilderness - Deschutes NF, Custer-Gallatin NF, Bridger-Teton NF, Lolo NF, Shasta-Trinity NF, Wells Gray Provincial Park, and Garibaldi Provincial Park. All other sites did not require official permission to conduct sampling.

We would like to thank Steffi Ickert-Bond at The University of Alaska Museum Herbarium (ALA); ALA accession numbers/Specimen GUID: V170482/UAM:Herb:244017; V153899/UAM:Herb:142478; V174721/UAM:Herb:251571; V174768/UAM:Herb:251929; V173773/UAM:Herb:250749).

---

**Figure S1:**
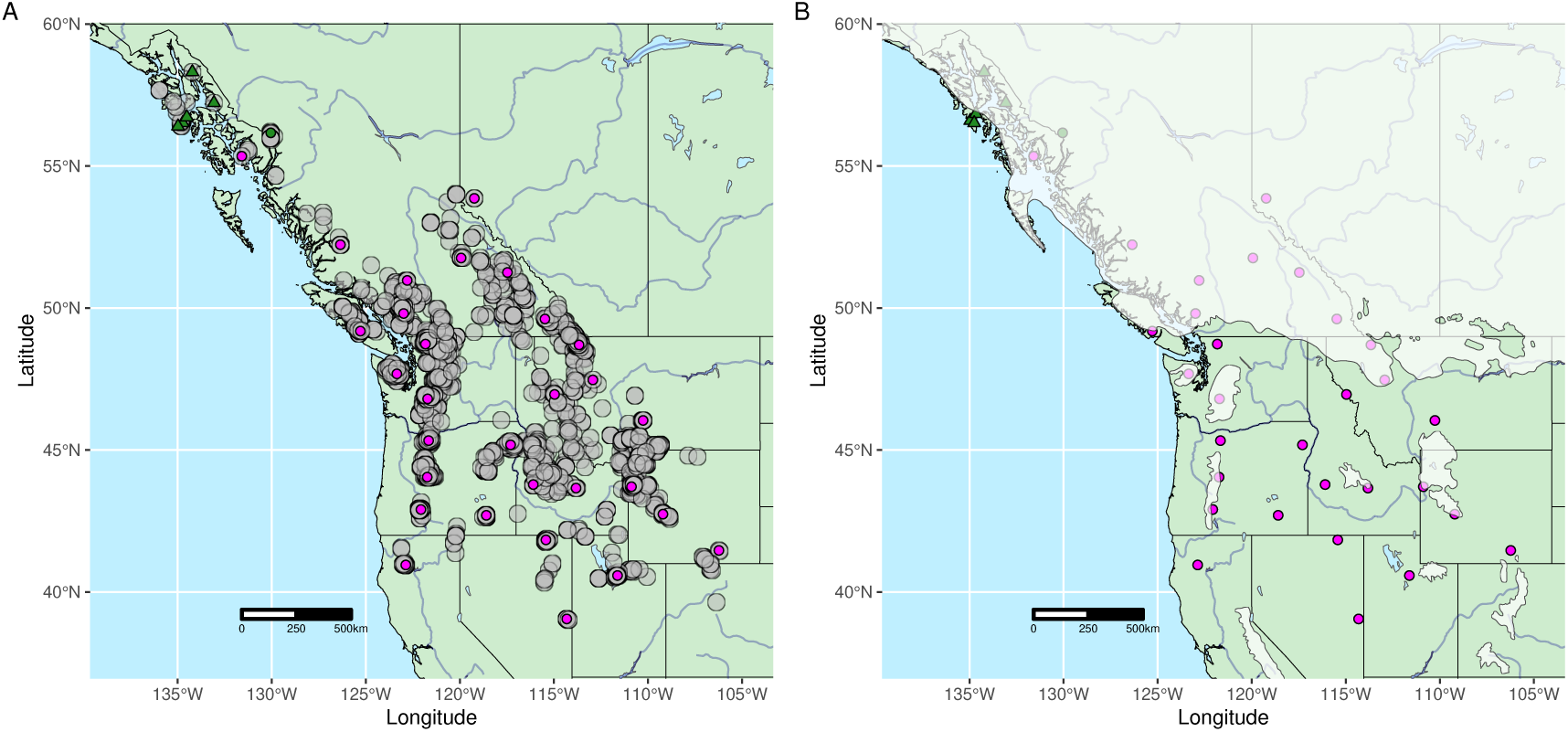
Sampling locations plotted with (A) range-wide occurrence records and (B) last glacial maximum of the Continental ice sheet.

**Figure S2:**
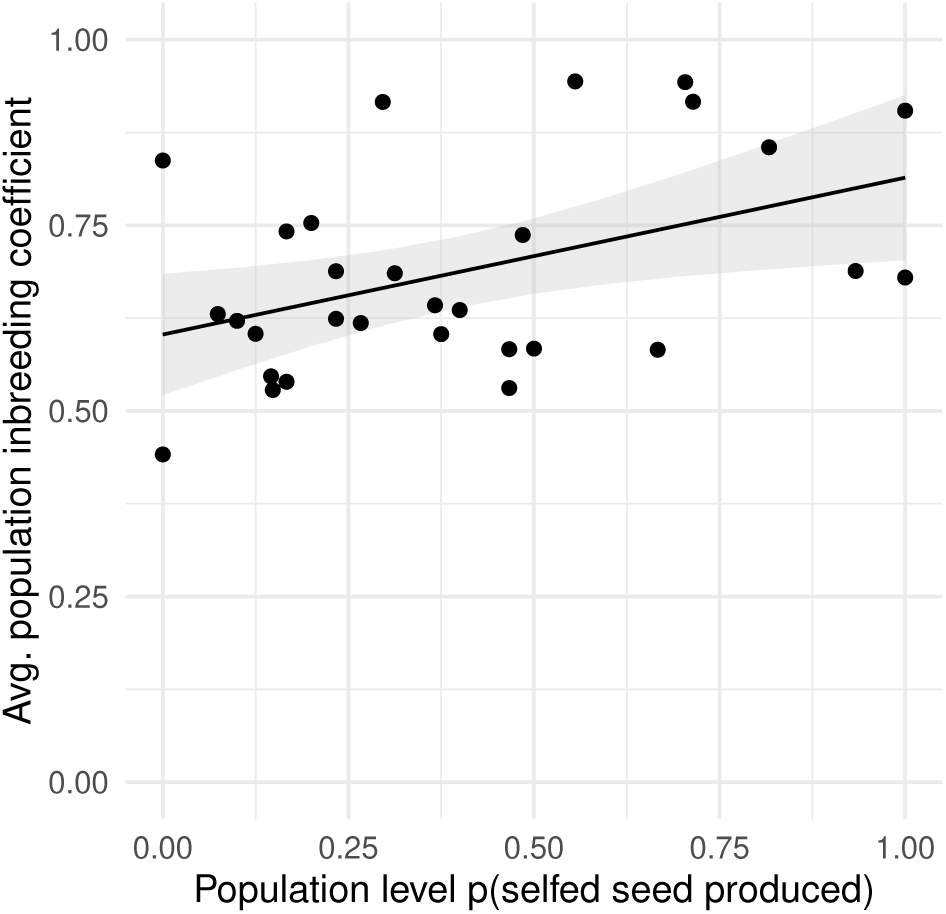
Population-level inbreeding coefficients (*F_IS_*) are positively correlated with population-level probability of selfing in the greenhouse.

**Figure S3:**
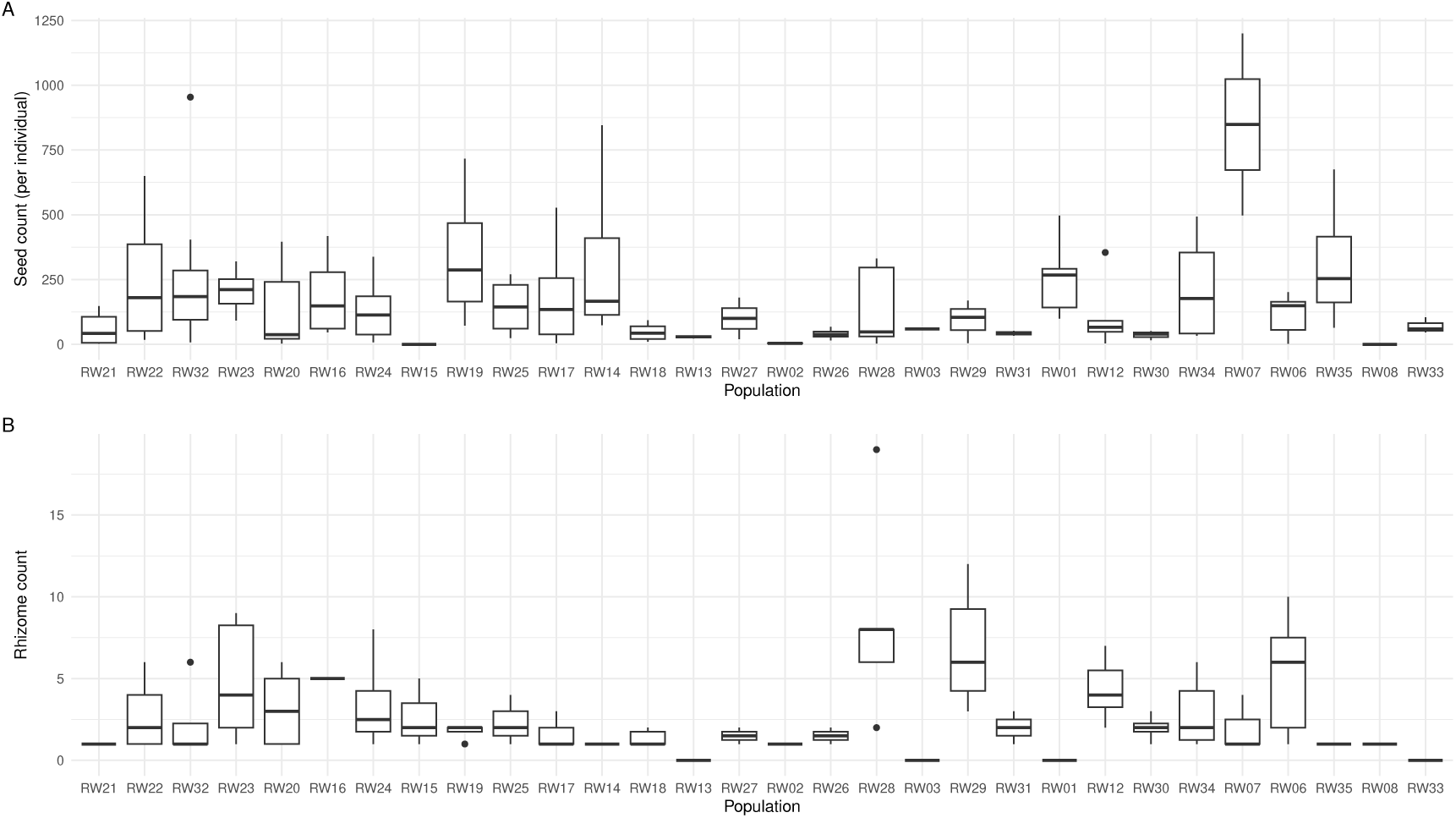
Boxplots of absolute seed count (A) and rhizome count (B) plotted by population, organized by increasing latitude.

**Figure S4:**
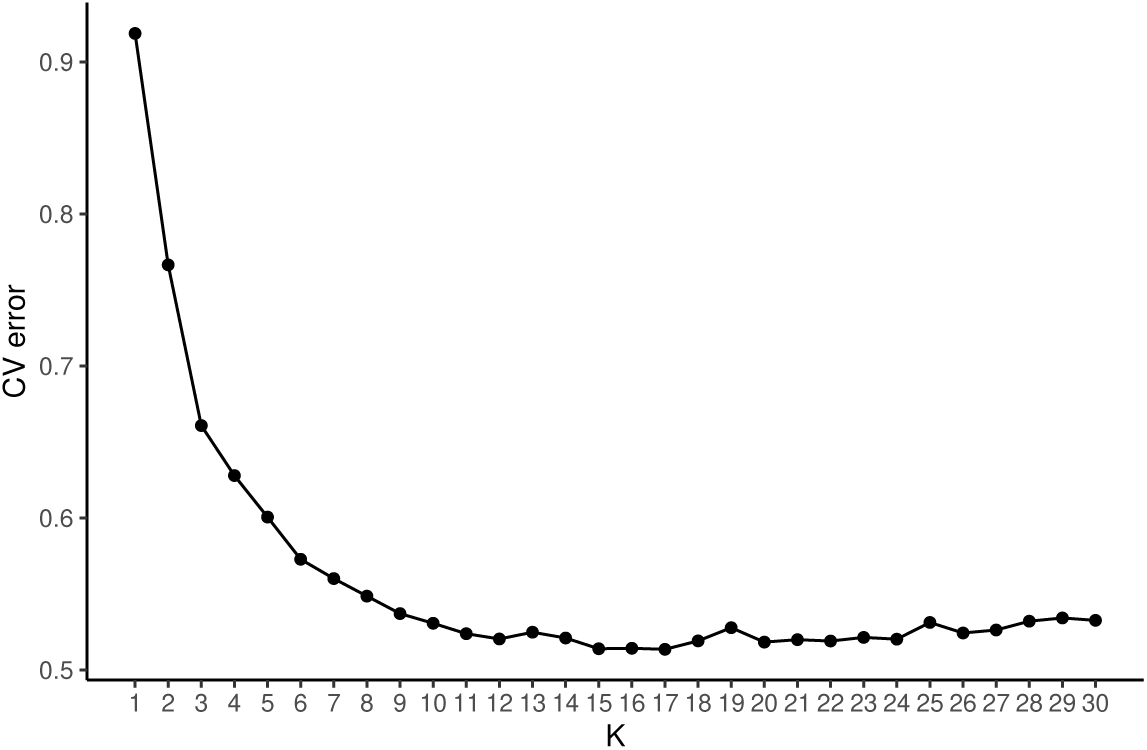
Cross-validation error plot from ADMIXTURE analysis.

**Figure S5:**
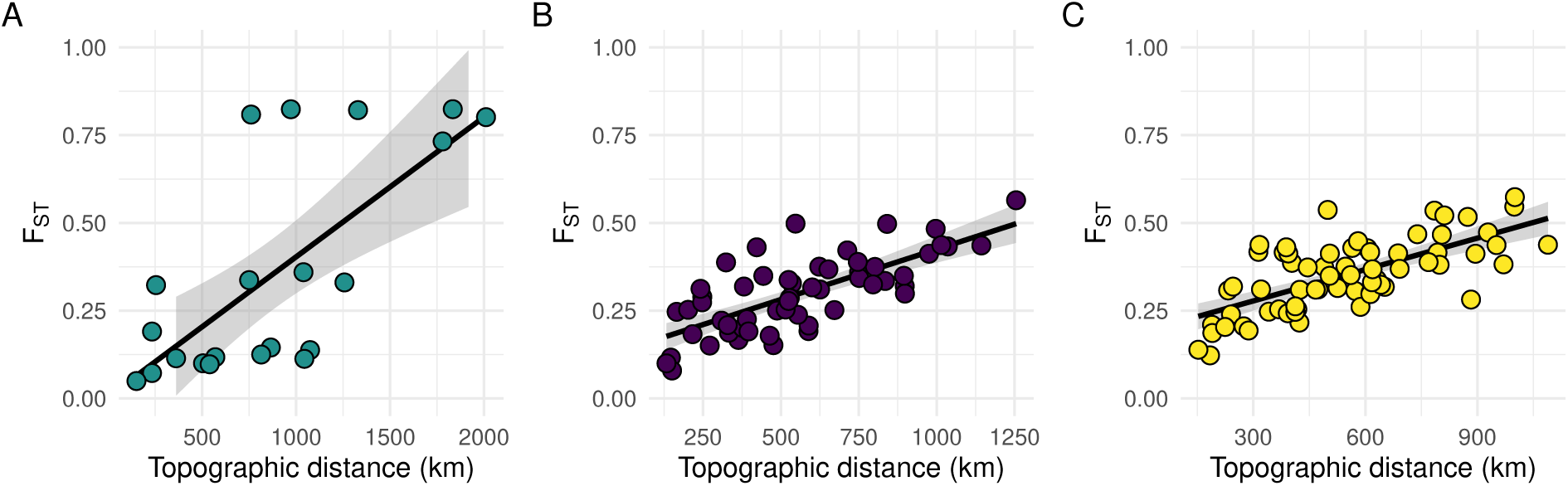
Significant signals of isolation-by-distance detected within each cluster. Mantel test A) Western cluster Mantel statistic *r* : 0.66, *p* = 0.008 B) Central cluster Mantel statistic *r* : 0.71, *p* < 0.0001 C) Eastern cluster Mantel statistic *r* : 0.63, *p* < 0.0001. For A), points around and above *F_ST_* ∼ 0.75 are comparisons with the population sampled in Northern California (RW32; Table 1). A significant IBD relationship in the Western cluster still exists even when RW32 is filtered out of the dataset (Mantel statistic *r* : 0.46, *p* = 0.04).

**Figure S6:**
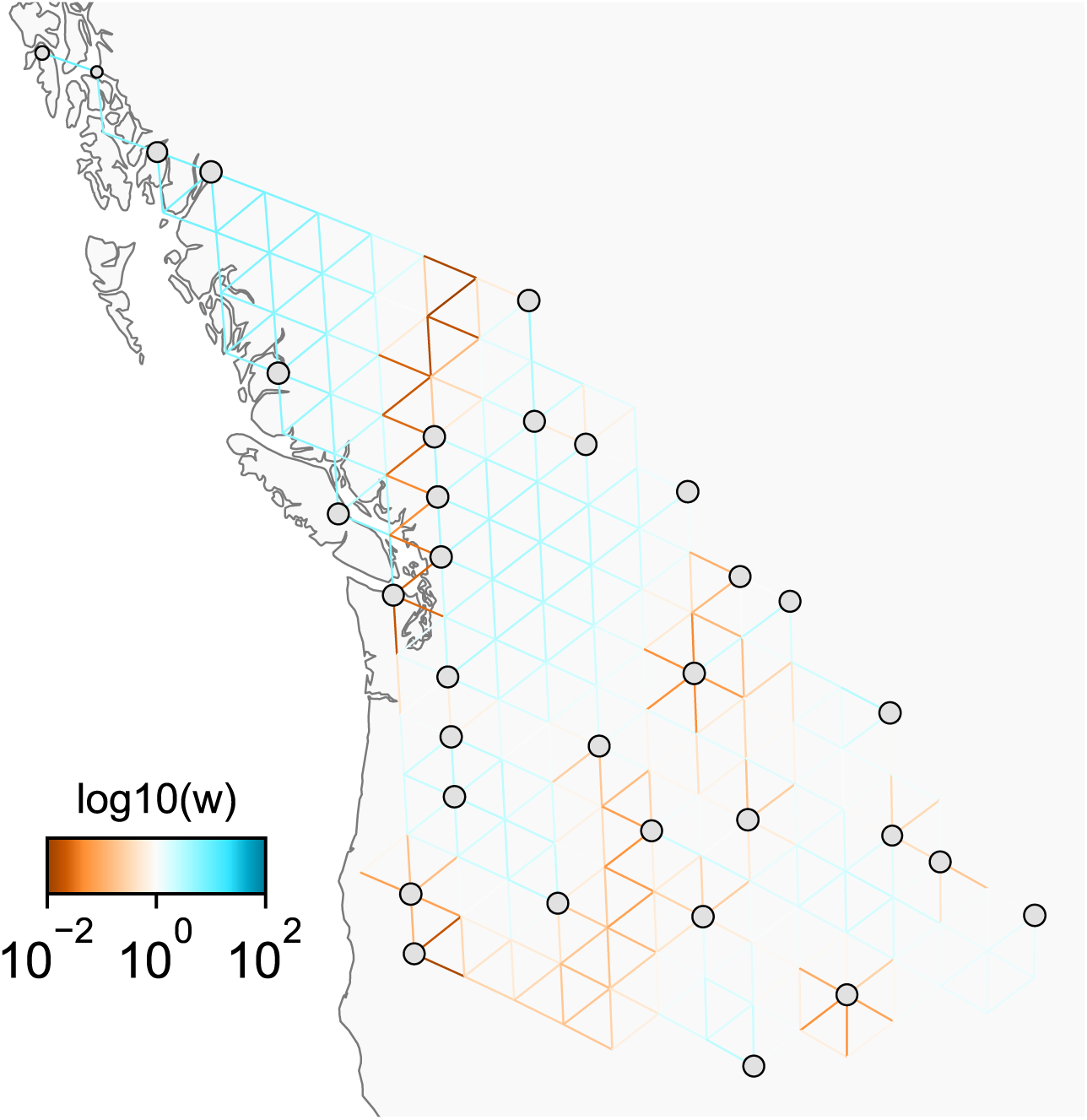
FEEMS analysis applied to the population genetic dataset generated from range wide sampling of *E. lewisii*. The grey points are the nodes assigned to the samples/sampling locations. In the inset legend, brown colours represent lower that average effective migration and blue colours represent higher than average effective migration. The migration surface reflects population structure and also geographic barriers (e.g., mountain ranges) that would prevent effective migration/gene flow.

**Figure S7:**
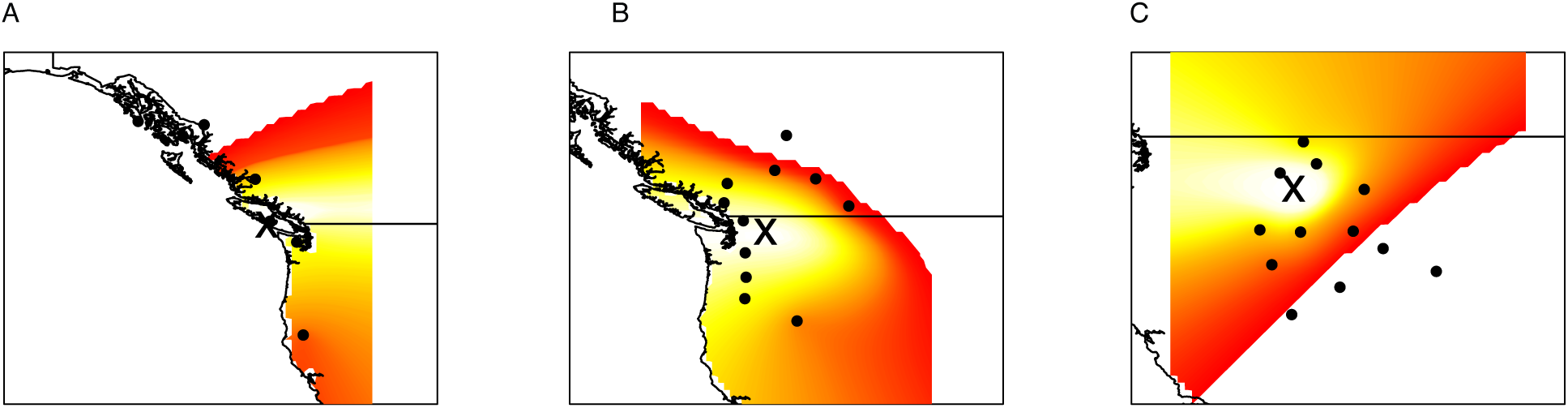
Locations of the most likely putative origin of range expansion within each ancestral cluster from the TDoA analysis (Peter and Slatkin, 2013) Location of putative historical refugia (marked by “*X*”) for A) Western (-125.2925, 49.3589), B) Central (-120.4843, 48.0656), and C) Eastern (-114.206, 46.0813) cluster. Heatmap indicates greater confidence in location of refugia in white-yellow (as measured by the sum of squares deviation) and low confidence in red areas (unlikely origin). Black points are sampling locations used to calculate the directionality index (*ψ*) and used to calculate pairwise geographic distances in TDoA analysis.

**Figure S8:**
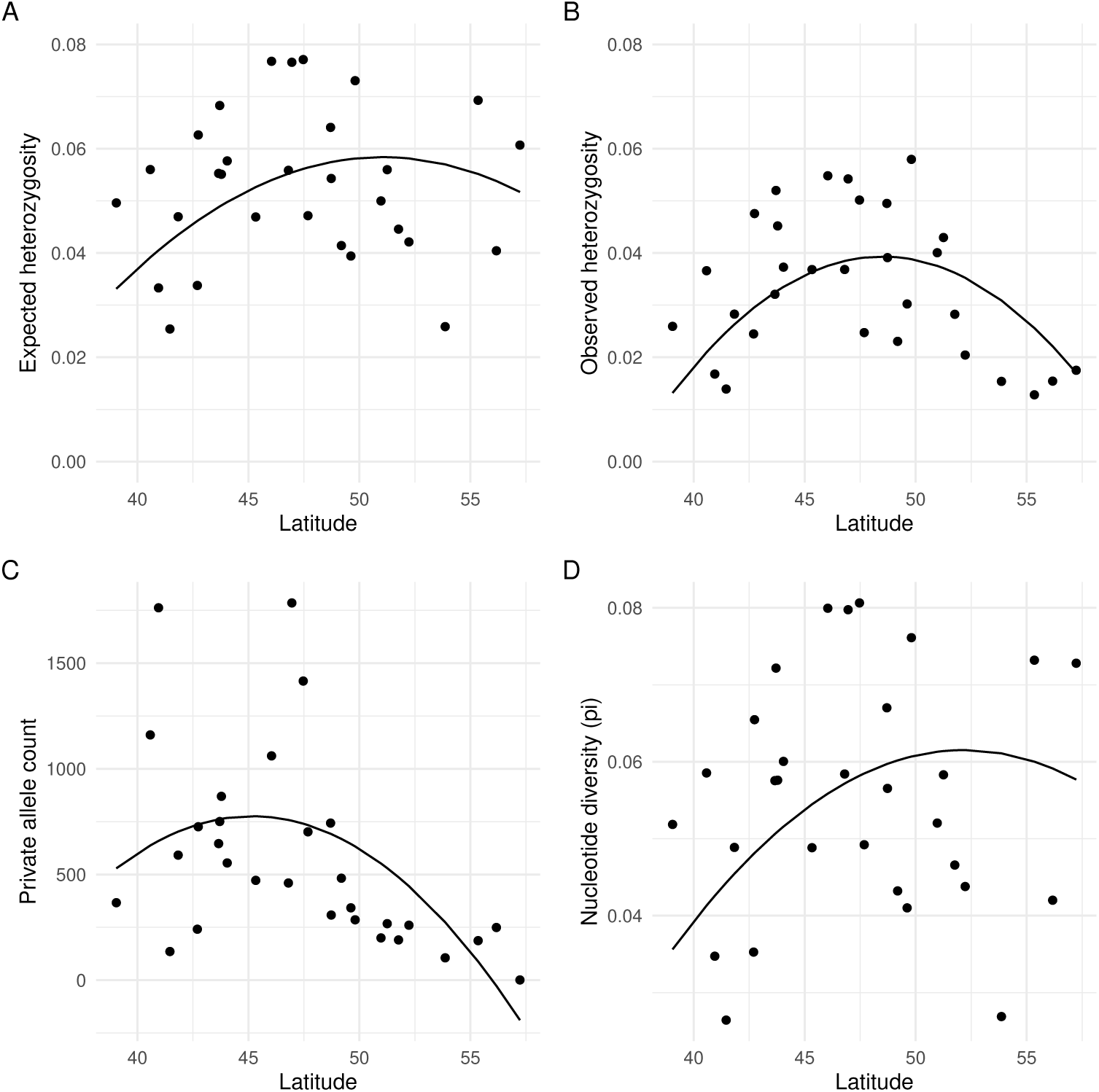
Plotting the best fit relationship between latitude and estimates of genetic diversity using quadratic (A–C) or linear (D) terms. Data points are population averages.

**Figure S9:**
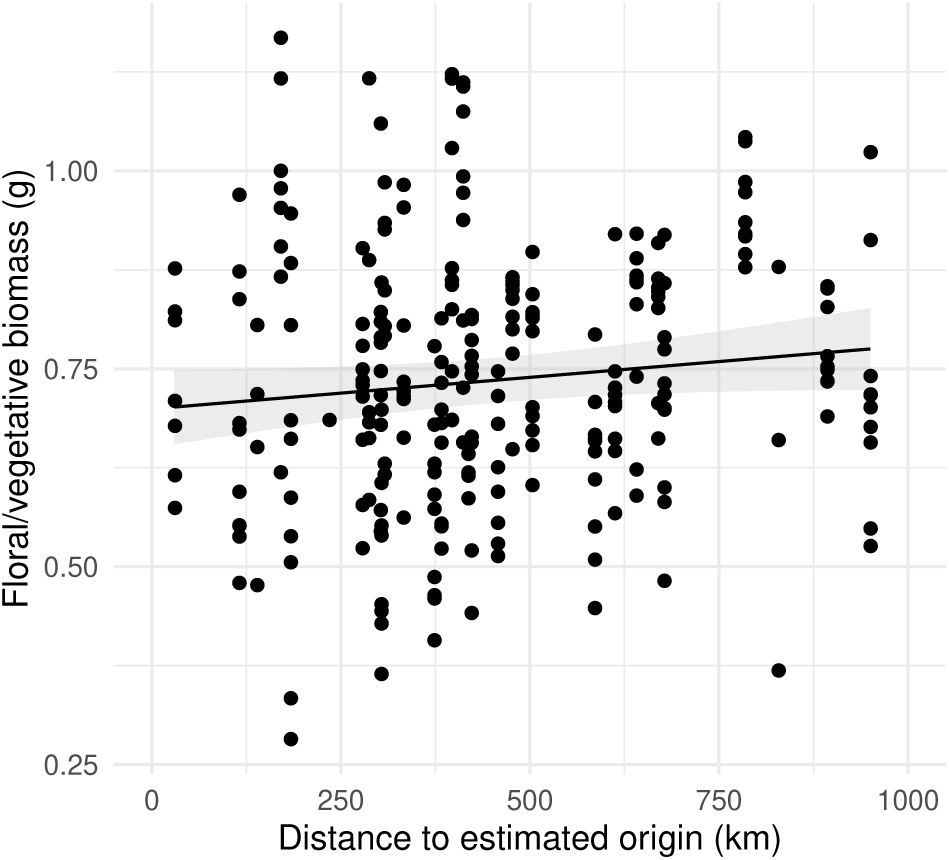
Average model output demonstrating that investment in floral tissues relative to vegetative tissues does not vary with distance from putative refugia. Data points are measurements from individual plants.

**Table S1:** Comparison of linear and quadratic model fit for the relationship between latitude and population-level estimates of genetic diversity.

| Response variable | linear model fit |  | quadratic model fit |  |
| --- | --- | --- | --- | --- |
| | $R^2$ | AIC | $R^2$ | AIC |
| Expected heterozygosity ($H_e$) | 0.44 | -142.86 | 0.48 | -143.56 |
| Observed heterozygosity ($H_o$) | 0.57 | -150.67 | 0.66 | -161.04 |
| Nucleotide diversity ($\pi$) | 0.44 | -139.24 | 0.47 | -139.03 |
| Private allele count | 0.29 | 433.60 | 0.5 | 410.86 |

## Literature Cited

1. Alexander, D. H., J. Novembre, and K. Lange. 2009. Fast model-based estimation of ancestry in unrelated individuals. Genome Research 19:1655–1664.

2. Austerlitz, F., B. Jung-Muller, B. Godelle, and P.-H. Gouyon. 1997. Evolution of Coalescence Times, Genetic Diversity and Structure during Colonization. Theoretical Population Biology 51:148–164.

3. Bai, W.-N., W.-J. Liao, and D.-Y. Zhang. 2010. Nuclear and chloroplast DNA phylogeography reveal two refuge areas with asymmetrical gene flow in a temperate walnut tree from East Asia. New Phytologist 188:892–901. eprint: https://onlinelibrary.wiley.com/doi/pdf/10.1111/j.1469-8137.2010.03407.x.

4. Baker, H. G. 1955. Self-compatibility and establishment after ”long distance” dispersal. Evolution 9:347–349.

5. Barrett, S. C., and L. D. Harder. 2017. The Ecology of Mating and Its Evolutionary Consequences in Seed Plants. Annual Review of Ecology, Evolution, and Systematics 48:135–157.

6. Barrett, S. C. H. 2015. Influences of clonality on plant sexual reproduction. Proceedings of the National Academy of Sciences 112:8859–8866. Publisher: Proceedings of the National Academy of Sciences.

7. Bates, D., M. Mächler, B. Bolker, and S. Walker. 2015. Fitting Linear Mixed-Effects Models Using lme4. Journal of Statistical Software 67:1–48.

8. Brown, J. H. 1984. On the Relationship between Abundance and Distribution of Species | The American Naturalist: Vol 124, No 2. The American Naturalist 124:255–279.

9. Busch, J. W. 2005. The evolution of self-compatibility in geographically peripheral populations of Leavenworthia alabamica (Brassicaceae). American Journal of Botany 92:1503–1512. eprint: https://onlinelibrary.wiley.com/doi/pdf/10.3732/ajb.92.9.1503.

10. Busch, J. W., and L. F. Delph. 2012. The relative importance of reproductive assurance and automatic selection as hypotheses for the evolution of self-fertilization. Annals of Botany 109:553–562.

11. Byers, D. L., and D. M. Waller. 1999. Do Plant Populations Purge Their Genetic Load? Effects of Population Size and Mating History on Inbreeding Depression. Annual Review of Ecology and Systematics 30:479–513. eprint: 10.1146/annurev.ecolsys.30.1.479.

12. Catchen, J., P. A. Hohenlohe, S. Bassham, A. Amores, and W. A. Cresko. 2013. Stacks: an analysis tool set for population genomics. Molecular Ecology 22:3124–3140. eprint: https://onlinelibrary.wiley.com/doi/pdf/10.1111/mec.12354.

13. Catchen, J. M., A. Amores, P. Hohenlohe, W. Cresko, and J. H. Postlethwait. 2011. Stacks: Building and Genotyping Loci De Novo From Short-Read Sequences. G3 Genes|Genomes|Genetics 1:171–182.

14. Charlesworth, D. 2006. Evolution of Plant Breeding Systems. Current Biology 16:R726–R735.

15. Charpentier, A. 2001. Consequences of clonal growth for plant mating. Evolutionary Ecology 15:521–530.

16. Cheptou, P.-O. 2012. Clarifying Baker’s Law. Annals of Botany 109:633–641.

17. Crnokrak, P., and S. C. H. Barrett. 2002. Purging the genetic load: a review of the experimental evidence. Evolution 56:2347–2358.

18. Dalton, A. S., M. Margold, C. R. Stokes, L. Tarasov, A. S. Dyke, R. S. Adams, S. Allard, H. E. Arends, N. Atkinson, J. W. Attig, P. J. Barnett, R. L. Barnett, M. Batterson, P. Bernatchez, H. W. Borns, A. Breckenridge, J. P. Briner, E. Brouard, J. E. Campbell, A. E. Carlson, J. J. Clague, B. B. Curry, R.-A. Daigneault, H. Dubé-Loubert, D. J. Easterbrook, D. A. Franzi, H. G. Friedrich, S. Funder, M. S. Gauthier, A. S. Gowan, K. L. Harris, B. Hétu, T. S. Hooyer, C. E. Jennings, M. D. Johnson, A. E. Kehew, S. E. Kelley, D. Kerr, E. L. King, K. K. Kjeldsen, A. R. Knaeble, P. Lajeunesse, T. R. Lakeman, M. Lamothe, P. Larson, M. Lavoie, H. M. Loope, T. V. Lowell, B. A. Lusardi, L. Manz, I. McMartin, F. C. Nixon, S. Occhietti, M. A. Parkhill, D. J. Piper, A. G. Pronk, P. J. Richard, J. C. Ridge, M. Ross, M. Roy, A. Seaman, J. Shaw, R. R. Stea, J. T. Teller, W. B. Thompson, L. H. Thorleifson, D. J. Utting, J. J. Veillette, B. C. Ward, T. K. Weddle, and H. E. Wright. 2020. An updated radiocarbon-based ice margin chronology for the last deglaciation of the North American Ice Sheet Complex. Quaternary Science Reviews 234:106223.

19. Danecek, P., J. K. Bonfield, J. Liddle, J. Marshall, V. Ohan, M. O. Pollard, A. Whitwham, T. Keane, S. A. McCarthy, R. M. Davies, and H. Li. 2021. Twelve years of SAMtools and BCFtools. GigaScience 10:giab008.

20. Dole, J. A. 1992. Reproductive Assurance Mechanisms in Three Taxa of the Mimulus guttatus Complex (Scrophulariaceae). American Journal of Botany 79:650–659. Publisher: Botanical Society of America.

21. Dray, S., and A.-B. Dufour. 2007. The ade4 Package: Implementing the Duality Diagram for Ecologists. Journal of Statistical Software 22:1–20.

22. Eckert, C. G. 2000. Contributions of Autogamy and Geitonogamy to Self-Fertilization in a Mass-Flowering, Clonal Plant. Ecology 81:532–542.

23. Fausto, J. A., V. M. Eckhart, and M. A. Geber. 2001. Reproductive assurance and the evolutionary ecology of self-pollination in *Clarkia xantiana* (Onagraceae). American Journal of Botany 88:1794–1800.

24. Ghazoul, J. 2005. Pollen and seed dispersal among dispersed plants. Biological Reviews 80:413– 443.

25. Goldberg, E. E., J. R. Kohn, R. Lande, K. A. Robertson, S. A. Smith, and B. Igic. 2010. Species Selection Maintains Self-Incompatibility. Science 330:493–495.

26. Goodwillie, C., R. D. Sargent, C. G. Eckert, E. Elle, M. A. Geber, M. O. Johnston, S. Kalisz, D. A. Moeller, R. H. Ree, M. Vallejo-Marin, and A. A. Winn. 2010. Correlated evolution of mating system and floral display traits in flowering plants and its implications for the distribution of mating system variation. New Phytologist 185:311–321.

27. Goudet, J., and T. Jombart. 2022. hierfstat: Estimation and Tests of Hierarchical F-Statistics.

28. Griffin, P. C., and Y. Willi. 2014. Evolutionary shifts to self-fertilisation restricted to geographic range margins in North American Arabidopsis lyrata. Ecology Letters 17:484–490. eprint: https://onlinelibrary.wiley.com/doi/pdf/10.1111/ele.12248.

29. Handel, S. N. 1985. The Intrusion of Clonal Growth Patterns on Plant Breeding Systems. The American Naturalist 125:367–384. Publisher: The University of Chicago Press.

30. Hargreaves, A. L., and C. G. Eckert. 2014. Evolution of dispersal and mating systems along geographic gradients: implications for shifting ranges. Functional Ecology 28:5–21. eprint: https://onlinelibrary.wiley.com/doi/pdf/10.1111/1365-2435.12170.

31. He, Q., J. R. Prado, and L. L. Knowles. 2017. Inferring the geographic origin of a range expansion: Latitudinal and longitudinal coordinates inferred from genomic data in an ABC framework with the program x - origin. Molecular Ecology 26:6908–6920.

32. Hemstrom, W., and M. Jones. 2023. snpR: User friendly population genomics for SNP data sets with categorical metadata. Molecular Ecology Resources 23:962–973. eprint: https://onlinelibrary.wiley.com/doi/pdf/10.1111/1755-0998.13721.

33. Herlihy, C. R., and C. G. Eckert. 2005. Evolution of self-fertilization at geographical range margins? A comparison of demographic, floral, and mating system variables in central vs. peripheral populations of Aquilegia canadensis (Ranunculaceae). American Journal of Botany 92:744–751. eprint: https://onlinelibrary.wiley.com/doi/pdf/10.3732/ajb.92.4.744.

34. Hewitt, G. 2000. The genetic legacy of the Quaternary ice ages. Nature 405:907–913. Publisher: Nature Publishing Group.

35. Hollister, J., T. Shah, J. Nowosad, A. L. Robitaille, M. W. Beck, and M. Johnson. 2023. elevatr: Access Elevation Data from Various APIs.

36. Igic, B., and J. R. Kohn. 2006. The distribution of plant mating systems: study bias against obligately outcrossing species. Evolution 60:1098–1103.

37. Kalisz, S., D. W. Vogler, and K. M. Hanley. 2004. Context-dependent autonomous self-fertilization yields reproductive assurance and mixed mating. Nature 430:884–887.

38. Kennedy, B. F., and E. Elle. 2008. The reproductive assurance benefit of selfing: importance of flower size and population size. Oecologia 155:469–477.

39. Kolis, K. M., C. S. Berg, T. C. Nelson, and L. Fishman. 2022. Population genomic consequences of life-history and mating system adaptation to a geothermal soil mosaic in yellow monkeyflowers. Evolution 76:765–781.

40. Korunes, K. L., and K. Samuk. 2021. pixy: Unbiased estimation of nucleotide diversity and divergence in the presence of missing data. Molecular Ecology Resources 21:1359–1368. eprint: https://onlinelibrary.wiley.com/doi/pdf/10.1111/1755-0998.13326.

41. Koski, M. H., D. L. Grossenbacher, J. W. Busch, and L. F. Galloway. 2017. A geographic cline in the ability to self-fertilize is unrelated to the pollination environment. Ecology 98:2930–2939.

42. Koski, M. H., N. C. Layman, C. J. Prior, J. W. Busch, and L. F. Galloway. 2019. Selfing ability and drift load evolve with range expansion. Evolution Letters 3:500–512.

43. Kuznetsova, A., P. B. Brockhoff, and R. H. B. Christensen. 2017. lmerTest Package: Tests in Linear Mixed Effects Models. Journal of Statistical Software 82. Publisher: Foundation for Open Access Statistic.

44. Li, H., and R. Durbin. 2009. Fast and accurate short read alignment with Burrows–Wheeler transform. Bioinformatics 25:1754–1760.

45. Li, H., B. Handsaker, A. Wysoker, T. Fennell, J. Ruan, N. Homer, G. Marth, G. Abecasis, R. Durbin, and 1000 Genome Project Data Processing Subgroup. 2009. The Sequence Alignment/Map format and SAMtools. Bioinformatics 25:2078–2079.

46. Marcus, J., W. Ha, R. F. Barber, and J. Novembre. 2021. Fast and flexible estimation of effective migration surfaces. eLife 10:e61927.

47. Marshall, D. L., J. J. Avritt, S. Maliakal-Witt, J. S. Medeiros, and M. G. M. Shaner. 2010. The impact of plant and flower age on mating patterns. Annals of Botany 105:7–22.

48. McKenna, A., M. Hanna, E. Banks, A. Sivachenko, K. Cibulskis, A. Kernytsky, K. Garimella, D. Altshuler, S. Gabriel, M. Daly, and M. A. DePristo. 2010. The Genome Analysis Toolkit: A MapReduce framework for analyzing next-generation DNA sequencing data. Genome Research 20:1297–1303.

49. McLachlan, J. S., J. S. Clark, and P. S. Manos. 2005. Molecular Indicators of Tree Migration Capacity Under Rapid Climate Change. Ecology 86:2088–2098. eprint: https://onlinelibrary.wiley.com/doi/pdf/10.1890/04-1036.

50. Oksanen, J., G. L. Simpson, G. Blanchet, R. Kindt, P. Legendre, P. R. Minchin, R. O’Hara, P. Solymos, M. H. H. Stevens, E. Szoecs, H. Wagner, M. Barbour, M. Bedward, B. Bolker, D. Borcard, G. Carvalho, M. Chirico, M. . Caceres}, S. Durand, H. B. A. Evangelista, R. FitzJohn, M. Friendly, B. Furneaux, G. Hannigan, M. O. Hill, L. Lahti, D. McGlinn, M.-H. Ouellette, E. . Cunha, T. Smith, A. Stier, C. J. . Braak, and J. Weedon. 2022. vegan: Community Ecology Package.

51. Opedal, O. H., E. Albertsen, W. S. Armbruster, R. Pérez-Barrales, M. Falahati-Anbaran, and C. Pélabon. 2016. Evolutionary consequences of ecological factors: pollinator reliability predicts mating-system traits of a perennial plant. Ecology Letters 19:1486–1495.

52. Pannell, J. R., J. R. Auld, Y. Brandvain, M. Burd, J. W. Busch, P.-O. Cheptou, J. K. Conner, E. E. Goldberg, A.-G. Grant, D. L. Grossenbacher, S. M. Hovick, B. Igic, S. Kalisz, T. Petanidou, A. M. Randle, R. R. de Casas, A. Pauw, J. C. Vamosi, and A. A. Winn. 2015. The scope of Baker’s law. New Phytologist 208:656–667.

53. Peischl, S., I. Dupanloup, M. Kirkpatrick, and L. Excoffier. 2013. On the accumulation of deleterious mutations during range expansions. Molecular Ecology 22:5972–5982.

54. Peischl, S., and L. Excoffier. 2015. Expansion load: recessive mutations and the role of standing genetic variation. Molecular Ecology 24:2084–2094.

55. Peter, B. M., and M. Slatkin. 2013. Detecting range expansions from genetic data: detecting range expansions from genetic data. Evolution 67:3274–3289.

56. Peterson, B. J., and W. R. Graves. 2016. Chloroplast phylogeography of Dirca palustris L. indicates populations near the glacial boundary at the Last Glacial Maximum in eastern North America. Journal of Biogeography 43:314–327. eprint: https://onlinelibrary.wiley.com/doi/pdf/10.1111/jbi.12621.

57. Petit, R. J., I. Aguinagalde, J.-L. De Beaulieu, C. Bittkau, S. Brewer, R. Cheddadi, R. Ennos, S. Fineschi, D. Grivet, M. Lascoux, A. Mohanty, G. Müller-Starck, B. Demesure-Musch, A. Palmé, J. P. Martín, S. Rendell, and G. G. Vendramin. 2003. Glacial Refugia: Hotspots But Not Melting Pots of Genetic Diversity. Science 300:1563–1565.

58. Petkova, D., J. Novembre, and M. Stephens. 2016. Visualizing spatial population structure with estimated effective migration surfaces. Nature genetics 48:94–100.

59. Pironon, S., G. Papuga, J. Villellas, A. L. Angert, M. B. García, and J. D. Thompson. 2017. Geographic variation in genetic and demographic performance: new insights from an old biogeographical paradigm. Biological Reviews 92:1877–1909. eprint: https://onlinelibrary.wiley.com/doi/pdf/10.1111/brv.12313.

60. Prior, C. J., and J. W. Busch. 2021. Selfing rate variation within species is unrelated to life-history traits or geographic range position. American Journal of Botany 108:2294–2308.

61. Pritchard, J. K., M. Stephens, and P. Donnelly. 2000. Inference of Population Structure Using Multilocus Genotype Data. Genetics 155:945–959.

62. Pujol, B., S.-R. Zhou, J. Sanchez Vilas, and J. R. Pannell. 2009. Reduced inbreeding depression after species range expansion. Proceedings of the National Academy of Sciences 106:15379– 15383.

63. Purcell, S., B. Neale, K. Todd-Brown, L. Thomas, M. A. Ferreira, D. Bender, J. Maller, P. Sklar, P. I. De Bakker, M. J. Daly, and P. C. Sham. 2007. PLINK: A Tool Set for Whole-Genome Association and Population-Based Linkage Analyses. The American Journal of Human Genetics 81:559–575.

64. Savary, P., J.-C. Foltete, H. Moal, G. Vuidel, and S. Garnier. 2021. graph4lg: A package for constructing and analysing graphs for landscape genetics in R. Methods in Ecology and Evolution 12:539–547. eprint: https://onlinelibrary.wiley.com/doi/pdf/10.1111/2041-210X.13530.

65. Schemske, D. W., and R. Lande. 1985. The evolution of self-fertilization and inbreeding depression in plants. ii. empirical observations. Evolution 39:41–52.

66. Shafer, A. B. A., C. I. Cullingham, S. D. Côté, and D. W. Coltman. 2010. Of glaciers and refugia: a decade of study sheds new light on the phylogeography of northwestern North America. Molecular Ecology 19:4589–4621.

67. Slatkin, M., and L. Excoffier. 2012. Serial Founder Effects During Range Expansion: A Spatial Analog of Genetic Drift. Genetics 191:171–181.

68. Vallejo-Marín, M., M. E. Dorken, and S. C. Barrett. 2010. The Ecological and Evolutionary Consequences of Clonality for Plant Mating. Annual Review of Ecology, Evolution, and Systematics 41:193–213.

69. Vaughton, G., and M. Ramsey. 2010. Pollinator-mediated selfing erodes the flexibility of the best-of-both-worlds mating strategy in Bulbine vagans. Functional Ecology 24:374–382. eprint: https://onlinelibrary.wiley.com/doi/pdf/10.1111/j.1365-2435.2009.01648.x.

70. Vickery, R. K., D. R. Phillips, and P. R. Wonsavage. 1986. Seed Dispersal in Mimulus guttatus by Wind and Deer. American Midland Naturalist 116:206.

71. Villegas, A. C. 2001. Spatial and Temporal Variability in Clonal Reproduction of Aechmea magdalenae, a Tropical Understory Herb1. Biotropica 33:48–59. eprint: https://onlinelibrary.wiley.com/doi/pdf/10.1111/j.1744-7429.2001.tb00156.x.

72. Waltari, E., R. J. Hijmans, A. T. Peterson, S. Nyári, S. L. Perkins, and R. P. Guralnick. 2007. Locating Pleistocene Refugia: Comparing Phylogeographic and Ecological Niche Model Predictions. PLOS ONE 2:e563. Publisher: Public Library of Science.

73. Wang, I. J. 2020. Topographic path analysis for modelling dispersal and functional connectivity: Calculating topographic distances using the topoDistance r package. Methods in Ecology and Evolution 11:265–272. eprint: https://onlinelibrary.wiley.com/doi/pdf/10.1111/2041-210X.13317.

74. Waser, N. M., R. K. Vickery, and M. V. Price. 1982. Patterns of seed disperal and population differentiation in *Mimulus guttatus*. Evolution 36:753–761.

75. Whitehead, M. R., R. Lanfear, R. J. Mitchell, and J. D. Karron. 2018. Plant Mating Systems Often Vary Widely Among Populations. Frontiers in Ecology and Evolution 6:38.

76. Willi, Y., M. Fracassetti, S. Zoller, and J. Van Buskirk. 2018. Accumulation of Mutational Load at the Edges of a Species Range. Molecular Biology and Evolution 35:781–791.

77. Xin, Z., and J. Chen. 2012. A high throughput DNA extraction method with high yield and quality. Plant Methods 8:26.

78. Zeitler, L., C. Parisod, and K. J. Gilbert. 2023. Purging due to self-fertilization does not prevent accumulation of expansion load. PLOS Genetics 19:e1010883.

